# Gain of Function Cytolysin Variant Expressed by Clinically Isolated Urinary *Enterococcus faecalis*

**DOI:** 10.64898/2026.10.02.756073

**Authors:** Neha V. Hulyalkar, Ibrahim O. Salaudeen, Humera Gull, Irene N. Hau, Namrata V. Sawant, Philippe E. Zimmern, Gabriele Meloni, Nicole J. De Nisco

## Abstract

Cytolysin is a toxin-bacteriocin commonly encoded by *Enterococcus faecalis* that can target erythrocytes, neutrophils, and macrophages as well as other Gram-positive bacteria. Cytolysin can lyse human, horse, rabbit, and mouse erythrocytes, but not sheep erythrocytes (SEs). It is hypothesized that SEs are resistant to cytolysin because they lack phosphatidylcholine (PC) in their membranes, while susceptible erythrocytes have PC-rich membranes, suggesting PC as a cytolysin binding target. Here, we isolated a urinary *E. faecalis* strain C33 that can efficiently lyse SEs and found that this activity was conserved among several *E. faecalis* clinical isolates. We demonstrated that although C33 cytolysin was necessary for SE lysis, it is not sufficient, suggesting a secondary factor enhances cytolysin activity against SEs. Through liposome assays we found that while PC was required for the activity of DS16 cytolysin, PC was dispensable for C33 cytolysin activity. To assess the biological significance of C33 cytolysin in the bladder microenvironment, we assessed C33 cytolysin activity against bladder epithelial cells and the urinary microbiota. We found that C33 cytolysin was more cytotoxic to human bladder epithelial cells than the canonical DS16 cytolysin. We also found greater inhibition of co-isolated urinary microbiota species *Streptococcus parasanguinis*, *Enterococcus raffinosus*, *Staphylococcus capitis*, and *Corynebacterium coyleae* by C33 cytolysin. Interestingly, *E. faecalis* C33 also inhibited the protective urogenital taxa *Lactobacillus crispatus* and *Bifidobacterium longum*. Together, these results reveal a new PC-independent cytolysin activity among clinical *E. faecalis* isolates that may support survival and competition in the urinary microenvironment.

**IMPORTANCE:** *Enterococcus faecalis* often causes hospital-acquired opportunistic infections, especially urinary tract infections. The virulence factor, cytolysin, is a plasmid-encoded toxin that can lyse red blood cells (RBCs), immune cells, and bacterial cells, and has been implicated in rabbit endocarditis and endophthalmitis models. *E. faecalis* cytolysin has been reported to lyse human, horse, rabbit, and mouse RBCs, but not sheep. This is thought to be due to the higher levels of phosphatidylcholine (PC) in susceptible RBCs, suggesting cytolysin targets PC. Here, we report a urinary *E. faecalis* strain C33 that lyses sheep RBCs and does not require PC for membrane lysis, unlike previously reported *E. faecalis* strains. Our findings suggest a potentially distinct target receptor for the gain of function C33 cytolysin. Additionally, we found enhanced C33 cytolysin activity against human bladder epithelial cells and the urinary microbiota, suggesting potentials roles in competition, colonization, or pathogenesis in the urinary niche.

## INTRODUCTION

*Enterococcus faecalis* is a Gram-positive facultative anaerobe commonly associated with the gastrointestinal tract of humans and animals (1). However, despite its commensal origins, *E. faecalis* has emerged as a significant public health concern, frequently associated with opportunistic hospital-acquired infections in susceptible hosts (2, 3). Intrinsic resistance to many antimicrobials coupled with acquired antibiotic resistance, has resulted in limited therapeutic options against enterococci (4). Over the past few decades, *E. faecalis* has emerged as an important cause of catheter-associated urinary tract infection (CAUTI), owing to its ability to adhere and form biofilms on a variety of surfaces (5–7).

Although *E. faecalis* is a gut commensal, its ability to readily acquire and transfer extrachromosomal virulence factors, facilitate invasion and colonization of the host (8). Most commonly encoded virulence factors include aggregation substance, surface adhesins, gelatinase, bacteriocins, and cytolysin (9). Cytolysin is a toxin and bacteriocin commonly expressed by various *E. faecalis* strains (10). This pore-forming toxin is encoded either on a conjugative, pheromone responsive pAD1 plasmid, or in the chromosome, and has been associated with increased toxicity in rabbit endophthalmitis and endocarditis models (10–14). The toxin is encoded by the cytolysin operon comprising of 5 biosynthetic genes (*cylL_L_*, *cylL_S_, cylM, cylB, cylA*) 1 immunity gene (*cylI*) and 2 divergently transcribed regulatory genes (*cylR2* & *cylR1*) (15–20). *E. faecalis* strains carrying cytolysin are reported to lyse a wide variety of cells like erythrocytes, neutrophils, macrophages, and other Gram-positive organisms (21). Previous studies have shown that human, horse, dog, rabbit, and mouse erythrocytes are susceptible to lysis by *E. faecalis* cytolysin (22). However, sheep erythrocytes were reported to be resistant to cytolysin (22). The differential susceptibility was attributed to higher levels of phosphatidylcholine (PC) in the erythrocyte membrane, hinting that PC might possibly be involved in binding to the toxin at the membrane surface (23, 24). However, little is known about the interaction between *E. faecalis* cytolysin and the target cell prior to lysis. Moreover, whether this toxin specifically binds to PC on the target cell remains elusive.

Here, we report clinical *E. faecalis* strains that have gained the ability to lyse sheep erythrocytes. We thoroughly characterize this phenotype in urinary *E. faecalis* isolate C33 and find that the pAD1-encoded cytolysin operon is necessary but not sufficient for lysis of sheep erythrocytes. We performed liposome leakage assays with various phospholipids to identify lipids required for cytolysin activity and found that while PC was required for canonical cytolysin activity, the gain of function C33 cytolysin activity does not require PC. Finally, to gain deeper understanding of implications of this gain of function cytolysin activity in *E. faecalis* competition and pathogenesis, we assessed activity against bladder epithelial cells and urinary microbiota species. C33 cytolysin demonstrated enhanced activity against bladder epithelial cells and various co-isolated urinary microbiota species.

Taken together, this study identifies a new *E. faecalis* cytolysin activity conserved amongst various clinical isolates that may confer a competitive advantage to *E. faecalis* within the urinary bladder environment.

## RESULTS

### Urinary *E. faecalis* C33 exhibits enhanced lysis of sheep erythrocytes

*E. faecalis* C33 was isolated from the urine of a postmenopausal woman with a history of recurrent urinary tract infection (UTI) as previously reported and was found to be hemolytic on sheep blood agar (**Table 1**) (25, 26). Quantitative hemolysis assays with *E. faecalis* C33 show hemolytic activity against both sheep erythrocytes (SEs) and rabbit erythrocytes (REs), while the cytolysin positive (cyl+) *E. faecalis* reference strain DS16 lysed only REs and not SEs (**FIG 1A, B**) (27, 28). Interestingly, C33 lysis of SEs but not REs was enhanced in anaerobic compared to ambient atmospheric conditions (**FIG 1A, B**). To further examine this phenotype, we quantified the hemolytic activity of C33 for 48 hours under four different atmospheric conditions ranging from ambient, 5% CO_2_, microaerophilic and anaerobic. We observed that C33 lysed REs and SEs in all the atmospheres, but C33 activity against SEs increased with a decrease in oxygen concentration (**FIG 1C, D**). Apart from DS16, we also compared the hemolytic activity of *E. faecalis* C33 with reference cyl+ MMH594 and cyl-negative OG1RF strains and observed that DS16 and MMH594 lysed REs as previously described (22) but unlike C33, did not lyse SEs (**FIG 1E**). Additionally, we compared the hemolytic activity of C33 with other urinary *E. faecalis* strains in our female urinary microbiome biobank that encode for cytolysin (**Table 1**) (26). We observed varying activity against SEs with 6 out of the 9 additional urinary strains exhibiting SE hemolytic activity (**FIG 1F**). Collectively, these results suggest that urinary *E. faecalis* C33 as well as 6 other urinary *E. faecalis* isolates display unique hemolytic activity against SEs unlike the previously reported *E. faecalis* cyl+ strains DS16 and MMH594.

**Figure 1.**
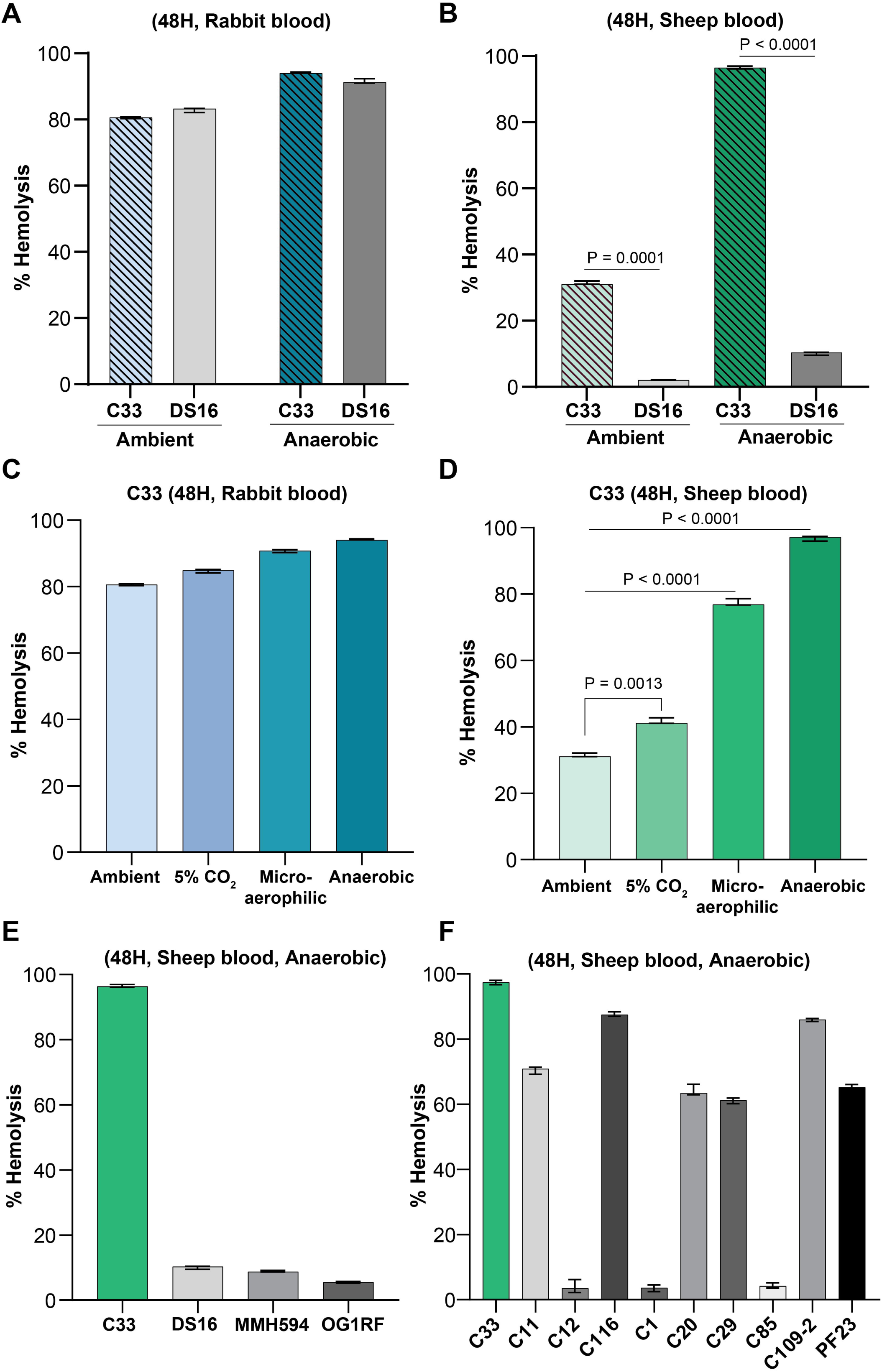
*E. faecalis* C33 is hemolytic against sheep erythrocytes. Quantitative hemolysis assays to compare hemolytic activity between (A) *E. faecalis* C33 & DS16 against rabbit erythrocytes (REs) and (B) against sheep erythrocytes (SEs); in ambient & anaerobic atmospheres after 48 hours. C33 activity against REs (C) and SEs (D) in ambient, 5% CO2, microaerophilic & anaerobic atmospheres. (E) Comparative quantitative hemolytic assays between *E. faecalis* C33 & reference strains DS16, MMH594 & OG1RF against SEs in anaerobic atmosphere. (F) Comparative quantitative hemolytic assays between E. faecalis C33 & other urinary strains against SEs in anaerobic atmosphere. P-values generated via the Brown-Forsythe and Welch ANOVA test.

**Table 1.** Bacterial strains used in this study.

| Bacterial species | Source | Presence of *E. faecalis* cytolysin | Reference |
| --- | --- | --- | --- |
| *E. faecalis* C33 | Urine | Yes; pAD1 plasmid | (26, 43) |
| *E. faecalis* DS16 | Clinical | Yes; pAD1 plasmid | (27, 28) |
| *E. faecalis* MMH594 | Blood | Yes; Chromosome | (57) |
| *E. faecalis* OG1RF | Oral | No | (58, 59) |
| *E. faecalis* C11 | Urine | Yes; pAD1 plasmid | (26) |
| *E. faecalis* C12 | Urine | Yes; pAD1 plasmid | (26) |
| *E. faecalis* C116 | Urine | Yes; pAD1 plasmid | (26) |
| *E. faecalis* C1 | Urine | Yes; Chromosome | (26) |
| *E. faecalis* C20 | Urine | Yes; Chromosome | (26) |
| *E. faecalis* C29 | Urine | Yes; Chromosome | (26) |
| *E. faecalis* C85 | Urine | Yes; Chromosome | (26) |
| *E. faecalis* C109-2 | Urine | Yes; Chromosome | (26) |
| *E. faecalis* PF23 | Urine | Yes; Chromosome | (26) |
| *Streptococcus parasanguinis* C33 | Urine | No | (43) |
| *Enterococcus raffinosus* C33 | Urine | No | (60) |
| *Staphylococcus capitis* C33 | Urine | No | (43) |
| *Corynebacterium coyleae* C33 | Urine | No | (43) |
| *Lactobacillus crispatus* Lc1226 | Urine | No | (48) |
| *Lactobacillus crispatus* Lc1700 | Urine | No | (48) |
| *Bifidobacterium longum* 1649 | Urine | No | (43) |

### Urinary *E. faecalis* C33 cytolysin is necessary for activity against the sheep erythrocytes

Previous studies have reported that *E. faecalis* cytolysin can lyse rabbit, horse and human erythrocytes but cannot lyse sheep erythrocytes (22). Moreover, this toxin can be encoded either on a self-transmissible, pheromone-responsive pAD1 plasmid or in a pathogenicity island (PAI) in the chromosome (12, 13). Analysis of our published complete C33 genome (GCA_030020615.1) revealed a cytolysin operon encoded on a pAD1-lineage plasmid, similar to the pAD1-encoded cytolysin operon in *E. faecalis* reference strain DS16 (Data S1) (26). Using DS16:pAD1 as reference, we performed a comparative analysis of the closed C33:pAD1 and DS16:pAD1 plasmid sequences using the BLAST Ring Image Generator (BRIG) v0.95 (**FIG 2A**) and observed 50% identity between both the plasmids and a conserved cytolysin operon. Furthermore, alignments of C33 cytolysin and DS16 cytolysin operon sequences showed 100% nucleotide identity between the two operons (**FIG 2B**). Since cytolysin has been implicated in the lysis of REs, we wanted to determine if it was also required for the lysis of SEs observed in *E. faecalis* C33 (22). To test the role of C33 cytolysin in SE lysis, we knocked-out the *cyl* operon by markerless gene exchange approach (29). On performing quantitative and qualitative hemolysis assays, we observed complete loss of SE and RE hemolysis in the C33Δ*cyl* mutant compared to the wild-type C33 strain (**FIG 2C, FIG S1A**). Complementing the C33Δ*cyl* mutant with *cyl* operon restored its hemolytic activity to wild-type levels against SEs and REs (**FIG 2C**, **FIG S1A**).

**Figure 2.**
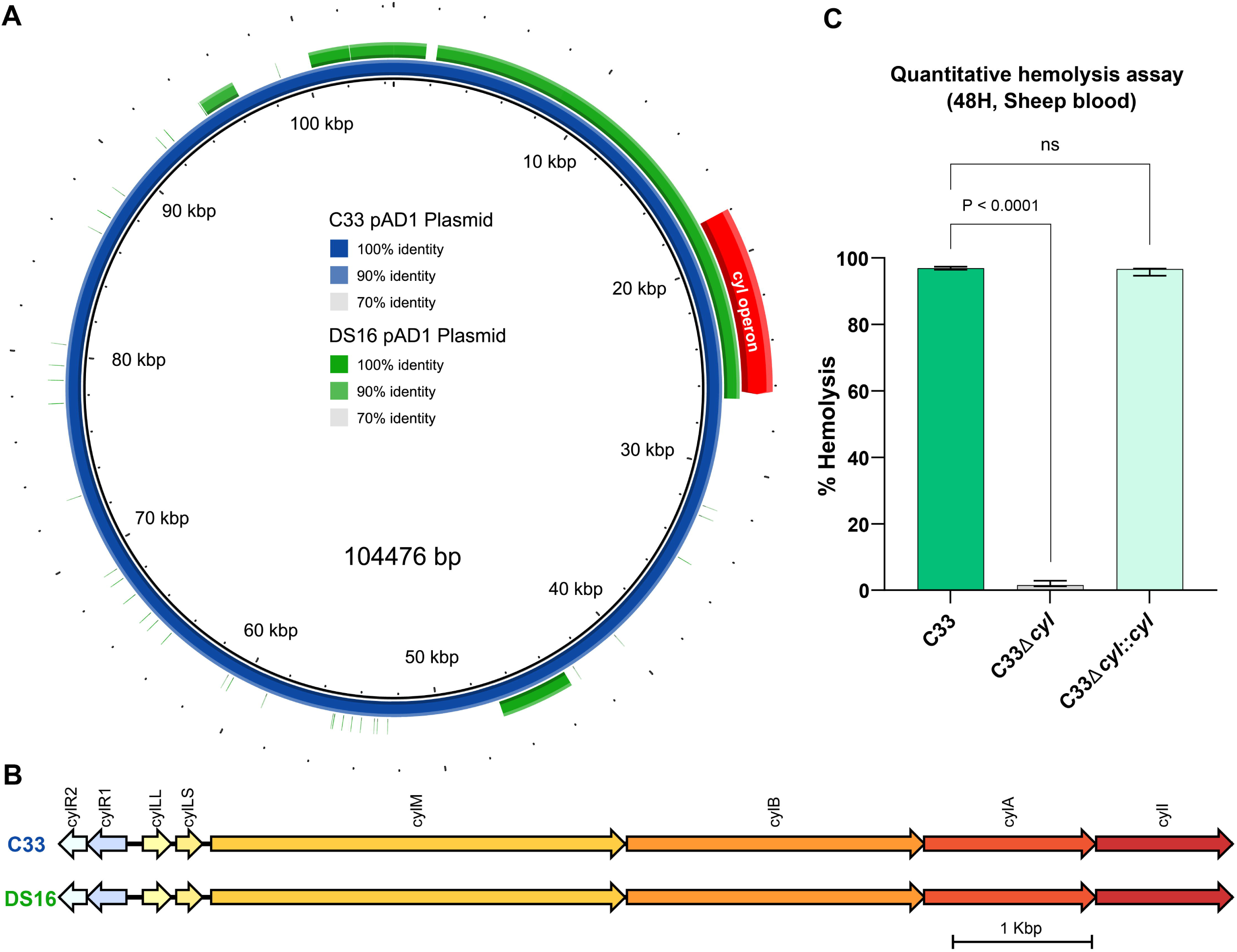
*E. faecalis* C33 lysis of sheep erythrocytes is cytolysin-dependent. **(A)** BLAST Atlas of pAD1 plasmids from *E. faecalis* C33 and DS16 constructed using pAD1 from C33 as a reference. **(B)** Blastn alignments of C33 cytolysin and DS16 cytolysin operons. **(C)** Quantitative hemolysis assays to test activity of C33, C33Δ*cyl* & C33Δ*cyl*::*cyl* against sheep erythrocytes. P-values generated via the Brown-Forsythe and Welch ANOVA test.

Since the C33 cytolysin operon was 100% identical to that of DS16, and knowing DS16 cannot lyse SEs, we wanted to test if this C33 enhanced hemolysis was due to differential expression of the cytolysin genes. We compared the expression of cytolysin operon genes by qPCR between C33, reference strains DS16, MMH594, and other urinary strains C11, C12, C116, C1, C20, C29, C85, C109-2 and PF23 in the presence of sheep blood and did not observe any significant difference in expression for any gene cyl operon genes between the strains (**FIG S1B-F**). Additionally, we also compared the expression of cytolysin genes in *E. faecalis* C33 between different atmospheres and did not observe any significant difference (**FIG S2**). Taken together, these results suggest that C33 *cyl* operon is necessary for hemolysis rabbit and sheep erythrocytes but is not significantly differentially expressed between isolates or in different atmospheres.

### The pAD1-encoded *E. faecalis* C33 cytolysin operon is not sufficient for lysis of sheep erythrocytes

Based on our finding that the C33 cytolysin operon was necessary for activity against sheep (SEs) erythrocytes (**FIG 2C**), we sought to determine if it was also sufficient. The *E. faecalis* C33 is encoded on a pAD1 plasmid that is a cAD1 pheromone-responsive and self-transmissible (30). Since OG1RF does not encode the cyl operon but produces the cAD1 pheromone, we mated the C33 pAD1 plasmid into OG1RF to begin to test for sufficiency of the C33 cytolysin operon in SE lysis. Prior to conjugation, the recipient OG1RF was transformed with a pCIE plasmid expressing C33 *cylA* and *cylI* to provide cytolysin immunity during mating. The transconjugant OG1RF:pAD1/2 expressed cytolysin genes when grown with rabbit or sheep blood, suggesting successful transfer of pAD1 plasmid to OG1RF (**FIG 3A**, **FIG S3A-G**). Interestingly, we also observed expression of *bacL1,* which is encoded by pAD2 plasmid, in the transconjugant, suggesting transfer of both pAD1 & pAD2 plasmids (**FIG 3B**). Moreover, the transconjugant was tested for expression of OG1RF *rhaB* gene, which is absent in *E. faecalis* C33, by qPCR to confirm its OG1RF parent strain identity (**FIG S3H, I**). On performing quantitative hemolysis assays, we observed that OG1RF:pAD1/2 lysed REs similar to C33 (**FIG 3C**) showing the cyl operon was not only expressed but functional in the transconjugant. However, OG1RF:pAD1/2 did not lyse SEs (**FIG 3D**), suggesting that the C33 pAD1 cytolysin operon is necessary but not sufficient for SE lysis.

**Figure 3.**
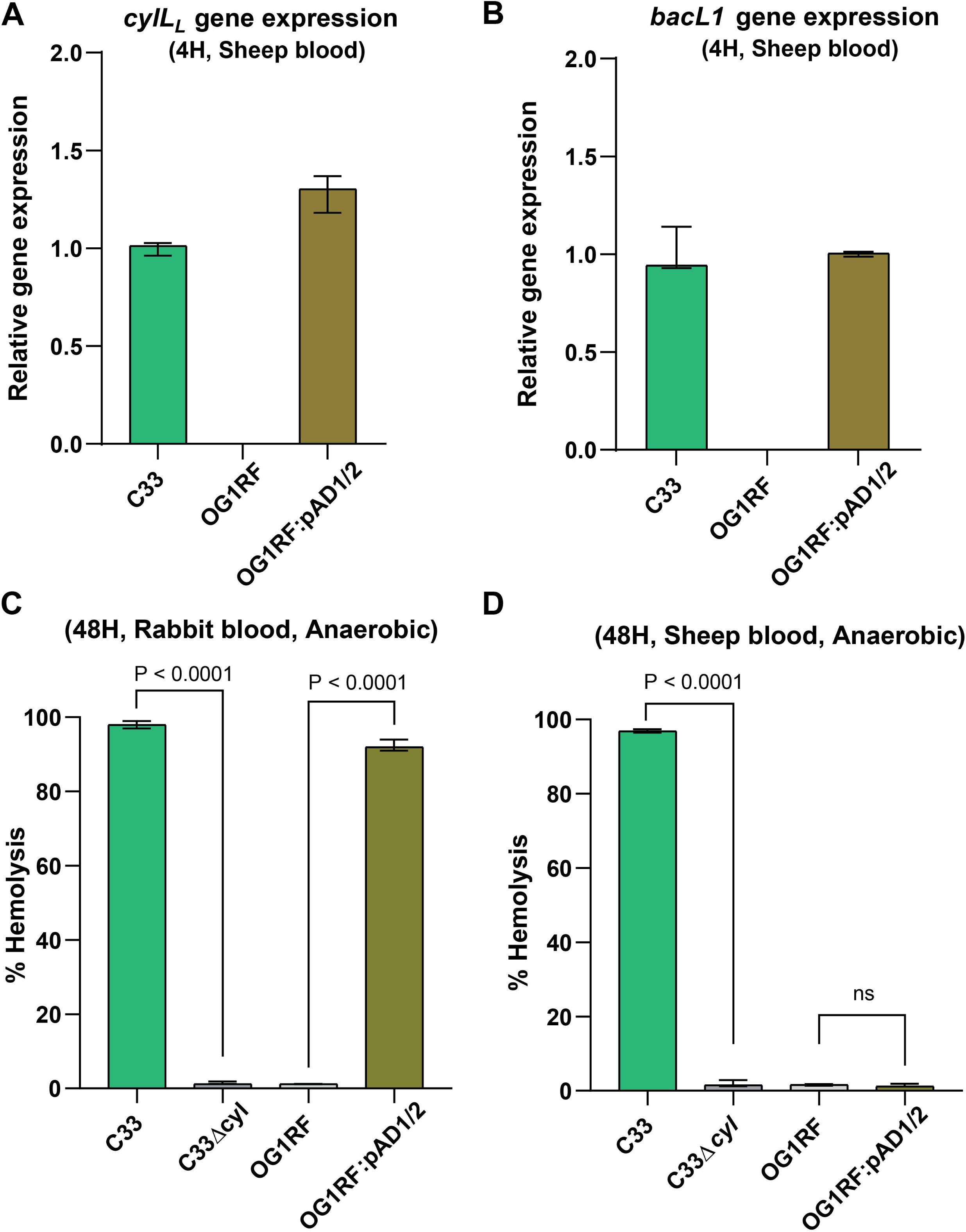
*E. faecalis* C33 pAD1 & pAD2 plasmids are insufficient for lysis of sheep erythrocytes. Quantitative real-time PCR to test expression of **(A)** *cylLL* gene (pAD1) and **(B)** *bacL1* gene (pAD2) in OG1RF:pAD1/2 transconjugant in the presence of sheep blood. Quantitative hemolysis assay to test the hemolytic activity of OG1RF:pAD1/2 transconjugant in **(C)** rabbit blood and **(D)** sheep blood. C33, C33Δ*cyl*, OG1RF were incorporated as positive and negative controls. P-values generated via the Brown-Forsythe and Welch ANOVA test.

### C33 cytolysin does not require phosphatidyl choline for membrane lysis unlike DS16

Miyazaki *et al.* reported that only human, horse, cow, and rabbit erythrocytes were susceptible to lysis by *E. faecalis* cytolysin, unlike sheep and goat erythrocytes which were resistant to cytolysin induced lysis (22). Since sheep and goat erythrocytes lack phosphatidyl choline (PC), it was speculated that lysis by this toxin is dependent on the presence of PC in the outer leaflet of susceptible erythrocyte membranes and that PC might act as a binding target (23). Since C33 lysed both SEs and REs, in contrast to DS16 which lysed only REs, we hypothesized that C33 might be targeting a non-PC phospholipid to lyse the erythrocytes. To test the dependence of C33 cytolysin activity individual phospholipids, we first generated liposomes composed of phospholipids that were representative of rabbit and sheep erythrocytes, and conducted liposome leakage assays (23, 24) (**Table S1**). Liposome lysis was measured by quantifying the release of the fluorescent 5(6)-carboxyfluorescein dye after incubation with the bacterial cultures. We observed that *E. faecalis* C33 was able to lyse both ‘sheep’ and ‘rabbit’ erythrocyte-like liposomes, whereas *E. faecalis* DS16 could lyse only the ‘rabbit’ liposomes (**FIG 4A, B**). The *E. faecalis* C33Δ*cyl* mutant was unable to lyse either sheep or rabbit erythrocyte-like liposomes, suggesting that the cytolysin toxin is necessary for C33 lysis of liposomes of both compositions (**FIG 4A, B**).

**Figure 4.**
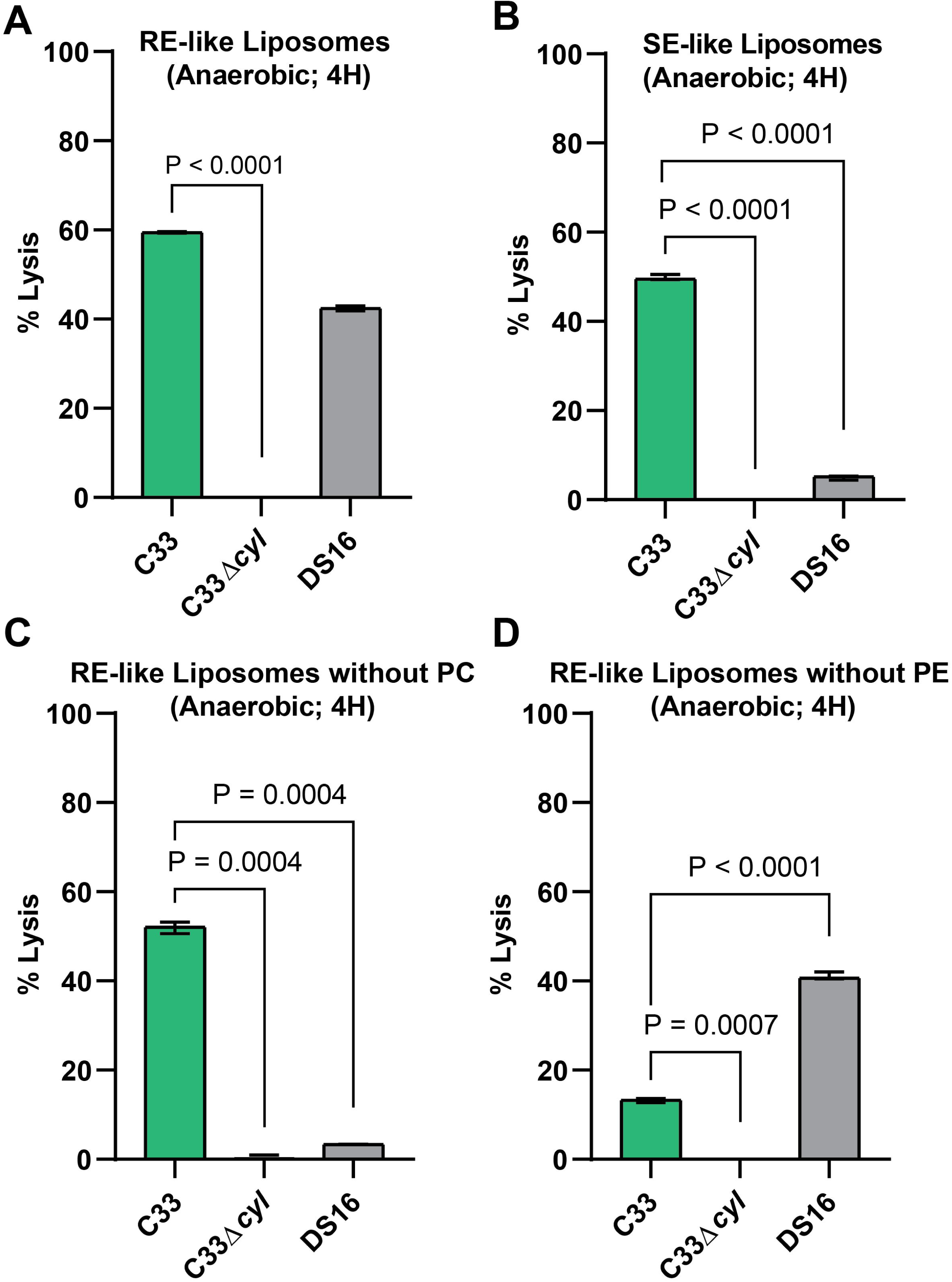
*E. faecalis* DS16 requires phosphatidyl choline for liposome lysis while *E. faecalis* C33 does not. Liposome leakage assay to test lytic activity of C33, C33Δ*cyl* & DS16 against **(A)** RE-like liposomes **(B)** SE-like liposomes **(C)** RE-like liposomes without phosphatidylcholine **(D)** RE-like liposomes without phosphatidylethanolamine. Composition of the liposomes is provided in Table 2. P-values generated via the Brown-Forsythe and Welch ANOVA test.

**Table 2.** Composition of the liposomes used in liposome leakage assay.

| Phospholipids | Rabbit Erythrocytes (RE) (%) | Sheep Erythrocytes (SE) (%) | RE-like w/o PC (%) | RE-like w/o PE (%) |
| --- | --- | --- | --- | --- |
| Phosphatidylethanolamine (PE) | 32 | 28 | 40 | 0 |
| Phosphatidylserine (PS) | 12 | 15 | 21 | 20 |
| Phosphatidylinositol (PI) | 2 | 3 | 11 | 10 |
| Phosphatidylcholine (PC) | 34 | 0 | 0 | 42 |
| Sphingomyelin (SM) | 20 | 54 | 28 | 28 |

To more directly assess the role of PC in cytolysin activity, we constructed liposomes of similar phospholipid composition of REs but removing PC (**Table S1**). These were termed as RE-like liposomes without PC. Removal of PC from the RE-like liposomes abolished *E. faecalis* DS16 cytolysin activity while *E. faecalis* C33 maintained lytic activity against the liposomes in a cytolysin dependent manner (**FIG 4C**). These data suggest that while PC is required for *E. faecalis* DS16 cytolysin activity, it is dispensable for C33 cytolysin activity. Because phosphatidylethanolamine (PE) is a major constituent of SEs, we hypothesized that C33 cytolysin activity may alternatively target PE (23). We compared the lytic activity of *E. faecalis* C33 and DS16 against RE-like liposomes without PE (**Table 2**). Interestingly, *E. faecalis* DS16 efficiently lysed RE-like liposomes without PE, likely due to the presence of PC in the liposome membrane (**FIG 4D**). Conversely, lysis by *E. faecalis* C33 was reduced in the absence of PE, even though PC was present, hinting at a possible role of PE in C33 gain-of-function cytolysin activity (**Figure 4D**). Importantly, the composition of sphingomyelin (SM), which also differs between REs and SEs, was constant between RE-like liposomes without PC or without PE, suggesting the observed differences in DS16 and C33 activity between the conditions were not due to sphingomyelin. Similar experiments were performed depleting other phospholipids found in REs and SEs, including SM, phosphatidylinositol (PI), and phosphatidylserine (PS) from RE-like liposomes (**Tabel S1**). However, no effect on *E. faecalis* DS16 or C33-mediated lysis was observed on depletion of SM, PI, or PS from RE-like liposomes (**FIG S4A-C**). Taken together, this data implicates PE in membrane lysis by *E. faecalis* C33 cytolysin activity unlike DS16, which requires PC for lysis.

### *E. faecalis* C33 cytolysin activity is strongly cytotoxic to bladder epithelial cells

Since gain of function cytolysin activity was observed in several urinary *E. faecalis* isolates, we postulated that it may have a functional role for *E. faecalis* fitness in the urinary environment. The bladder urothelium provides a strong barrier against bacterial infiltration; however, uropathogens like uropathogenic *Escherichia coli* utilize cytotoxins to lyse bladder epithelial cells and disrupt barrier function (31–33). As both PC and PE are present in bladder epithelial cell membranes, we hypothesized that *E. faecalis* C33 cytolysin activity would also efficiently lyse bladder epithelial cells (34–37). We therefore used a lactate dehydrogenase (LDH) release assay to assess *E. faecalis* C33 and DS16 cytotoxicity against 5637 bladder epithelial cells. The lysis of 5637 bladder epithelial cells infected with MOI 10 of *E. faecalis* strains was quantified 4 hours and 16 hours post-infection. *E. faecalis* C33 was significantly more cytotoxic to 5637 bladder epithelial cells than DS16 and this toxicity increased as time post-infection progressed **(FIG 5A, B**). C33 exhibited 80% toxicity against 5637 bladder epithelial cells after 16 hours, while DS16, with an 100% identical cytolysin operon as C33, exhibited only ∼25% cytotoxicity (**FIG 5B**). Moreover, the *E. faecalis* C33Δ*cyl* strain exhibited minimal (∼15%) cytotoxicity, suggesting that cytolysin is indeed necessary for the *E. faecalis* C33 cytotoxicity against cultured bladder epithelial cells (**FIG 5A, B**). These results indicate that the gain of function cytolysin activity of *E. faecalis* C33 strain enables enhanced cytotoxicity against urothelial cells, unlike *E. faecalis* DS16, suggesting a possible functional role in the urinary environment.

**Figure 5.**
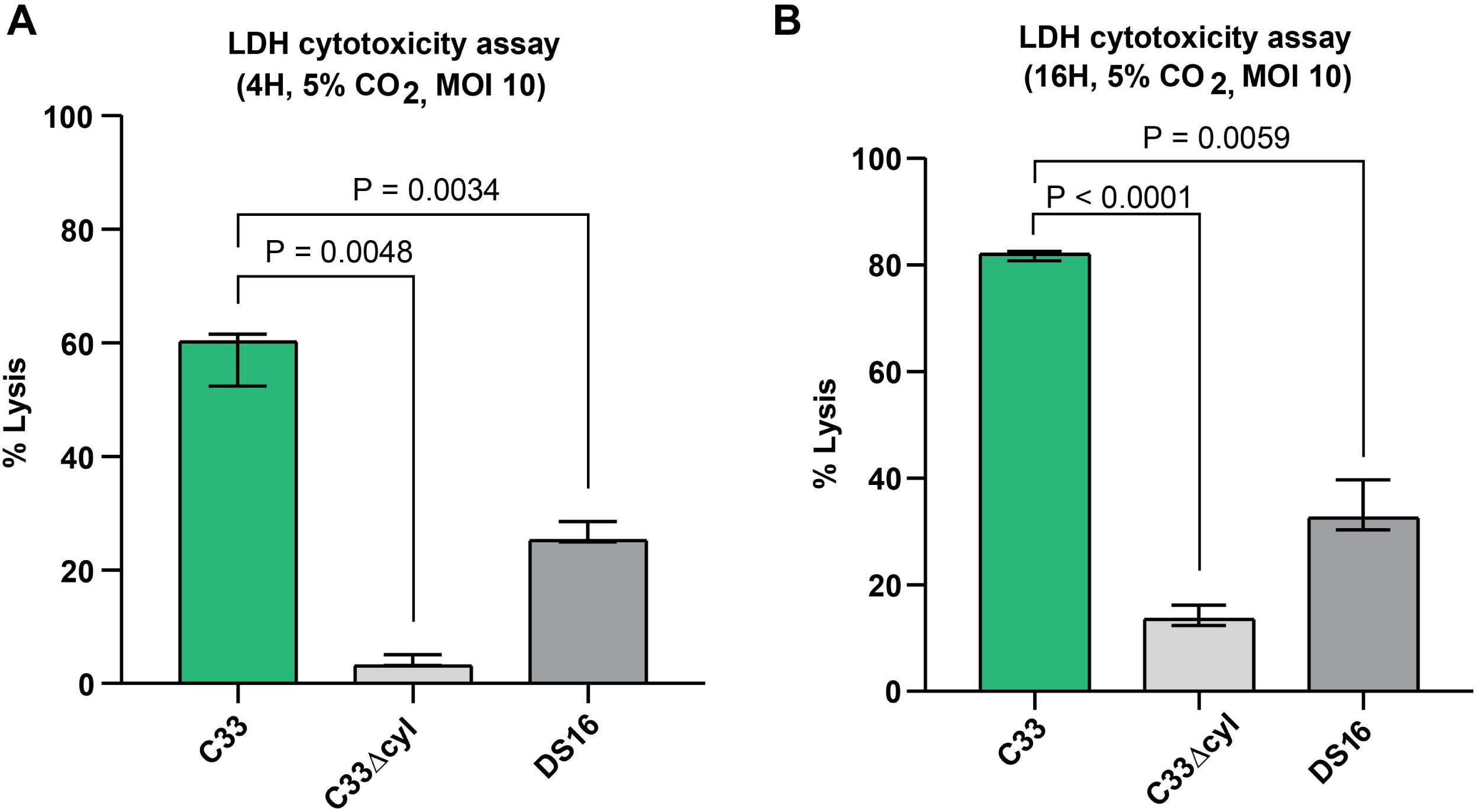
*E. faecalis* C33 lysed human bladder epithelial 5637 cells. LDH cytotoxicity assay to test *E. faecalis* C33 cytolysin activity against human bladder carcinoma 5637 cells at MOI 10 at **(A)** 4 hours and **(B)** 16 hours post-infection. P-values generated via the Brown-Forsythe and Welch ANOVA test.

### *E. faecalis* C33 cytolysin activity inhibits *E. faecalis* DS16 and OG1RF and is independent of pAD2-encoded Bac41

In addition to functioning as a cytotoxin against eukaryotic cells, *E. faecalis* cytolysin also acts as a bacteriocin against Gram-positive bacteria (38, 39). This bacteriocin activity is believed to provide a competitive advantage to cytolysin producing strains over non-cytolysin producers (10, 40). Hence, we hypothesized that urinary *E. faecalis* C33 with enhanced activity against eukaryotic cells, might have gained inhibitory activity against other cytolysin and non-cytolysin producing *E. faecalis* strains. To test this, we performed quantitative competition assays in BHI medium of C33 against *E. faecalis* reference strains DS16 and OG1RF. Interestingly, despite their 100% identical cytolysin operons, C33 inhibited DS16 (**FIG 6A**). However, the cytolysin deletion strain, C33Δ*cyl* did not inhibit DS16 (**FIG 6B**), implying that this gained activity was dependent upon the cytolysin operon. Both C33 and C33Δ*cyl* inhibited OG1RF (**FIG 6C, D**). On genome mining, we found bacteriocin Bac41 on the C33 pAD2 plasmid (41, 42) that could be inhibiting OG1RF (**FIG S5A**). We generated *E. faecalis* C33 *bacL1* single (C33Δ*bacL1*) as well as *cyl* and *bacL1* double (C33Δ*cyl*Δ*bacL1*) mutants and tested activity against OG1RF (**FIG S5B, C)**. We observed that although C33Δ*bacL1* still inhibited OG1RF, the double mutant strain, C33Δ*cyl*Δ*bacL1,* did not inhibit OG1RF (**FIG S5D, FIG 6C**). Thus, both C33 cytolysin and Bac41 bacteriocins may work together to inhibit OG1RF. To determine if Bac41 contributes to the C33 *E. faecalis* gain of function cytolysin activity against SEs, we measured SE hemolysis of the C33Δ*bacL1* single mutant and observed SE hemolysis levels similar to wildtype whereas no SE hemolysis was observed in the C33Δ*cyl* and C33Δ*cyl*Δ*bacL1* mutants (**FIG 6D**). The same trend was observed for RE hemolysis (**FIG S5E**). These data confirm that unlike the *cyl* operon, Bac41 is not necessary for SE lysis.

**Figure 6.**
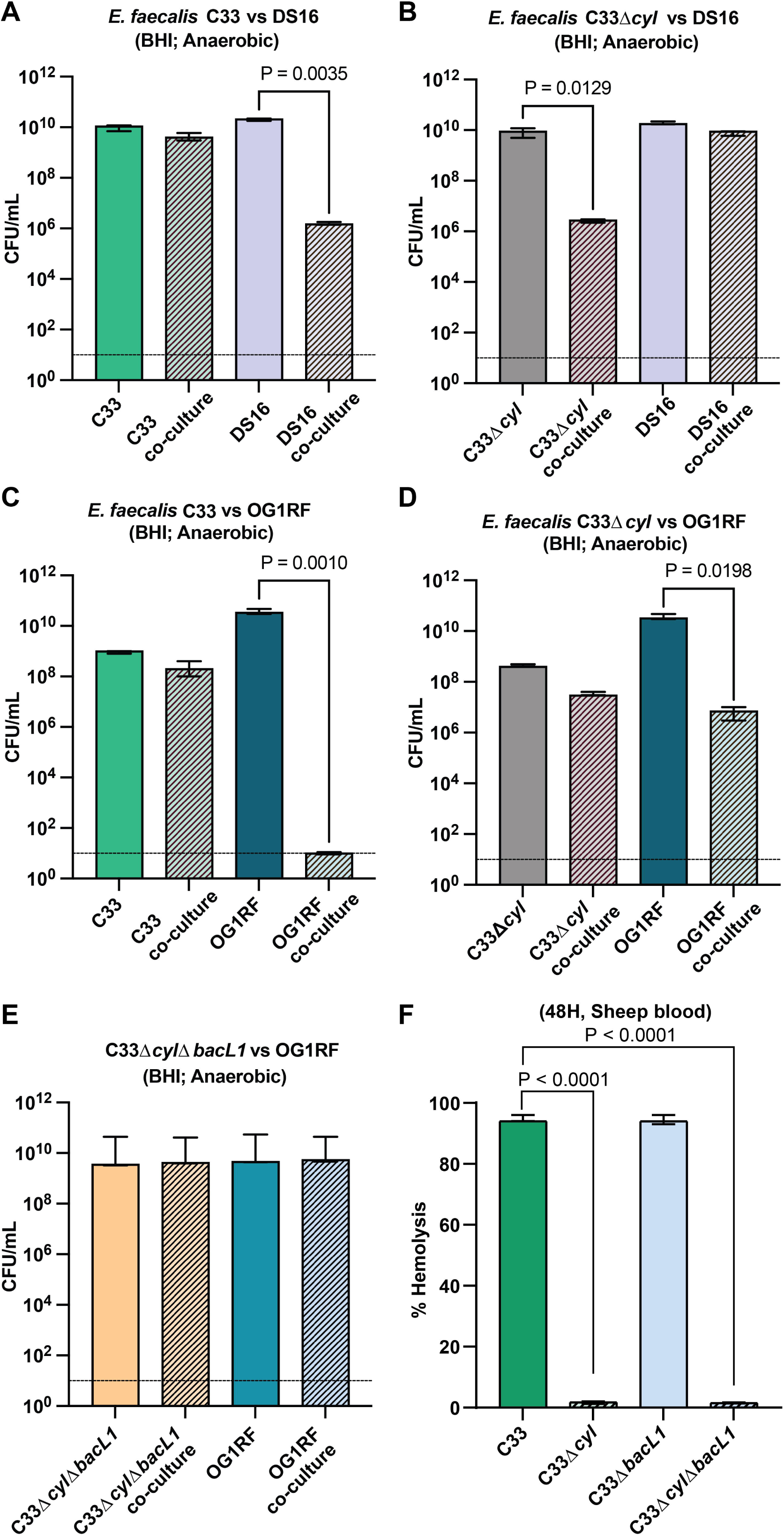
*E. faecalis* C33 cytolysin inhibits *E. faecalis* DS16 and OG1RF. Quantitative CFU competition assay to assess growth inhibition between **(A)** *E. faecalis* C33 WT and *E. faecalis* DS16 **(B)** *E. faecalis* C33Δ*cyl* and DS16, **(C)** *E. faecalis* C33 WT and OG1RF, **(D)** *E. faecalis* C33Δ*cyl* and OG1RF, and **(E)** *E. faecalis* C33Δ*bacL* and OG1RF. P-values were generated via Welch’s T-test. **(F)** Quantitative hemolysis assay to test the hemolytic activity of C33Δ*bacL* and C33Δ*cyl*Δ*bacL* against sheep erythrocytes. P-values were generated via the Brown-Forsythe and Welch ANOVA test.

### *E. faecalis* C33 cytolysin activity enhances competition against the urinary microbiota

Multiple bacterial species were co-isolated from the same urine sample as *E. faecalis* C33. These species included *Streptococcus parasanguinis, Enterococcus raffinosus, Staphylococcus capitis,* and *Corynebacterium coyleae* (43). We performed quantitative competition assays to test if C33 cytolysin would inhibit these co-isolated urinary microbiota species. For *S. parasanguinis* and *E. raffinosus,* assays were performed in Artificial Urine Medium (AUM^Y^) supplemented with 0.3% yeast extract (**Table S2**) (44, 45) under microaerophilic atmosphere. *E. faecalis* C33 significantly inhibited both *S. parasanguinis* and *E. raffinosus* in a cytolysin dependent manner and to a greater extent than *cyl+ E. faecalis* DS16 (**FIG 7A-F**). Indeed, *E. faecalis* C33 co-incubation resulted in ∼3 log(10) less *S. parasanguinis* and *E. raffinosus* CFUs than DS16 co-incubation (**FIG S6A, B**). We performed similar assays against co-isolated *S. capitis* and *C. coyleae* strains in Brain Heart Infusion (BHI) due to their inability to grow in AUM^Y^. We observed that *E. faecalis* C33 inhibited both *S. capitis* and *C. coyleae* in a cytolysin dependent manner and to a much greater extent than *E. faecalis* DS16 (**FIG 7G-L**). Incubation with *E. faecalis* C33 resulted in ∼3 log(10) less *S. capitis* and ∼2 log(10) less *C. coyleae* than did incubation with *E. faecalis* DS16 (**FIG S6C, D**). Taken together these results suggest that *E. faecalis* C33 gain of function cytolysin activity enhances *E. faecalis* competition against co-isolated urinary microbiota species.

**Figure 7.**
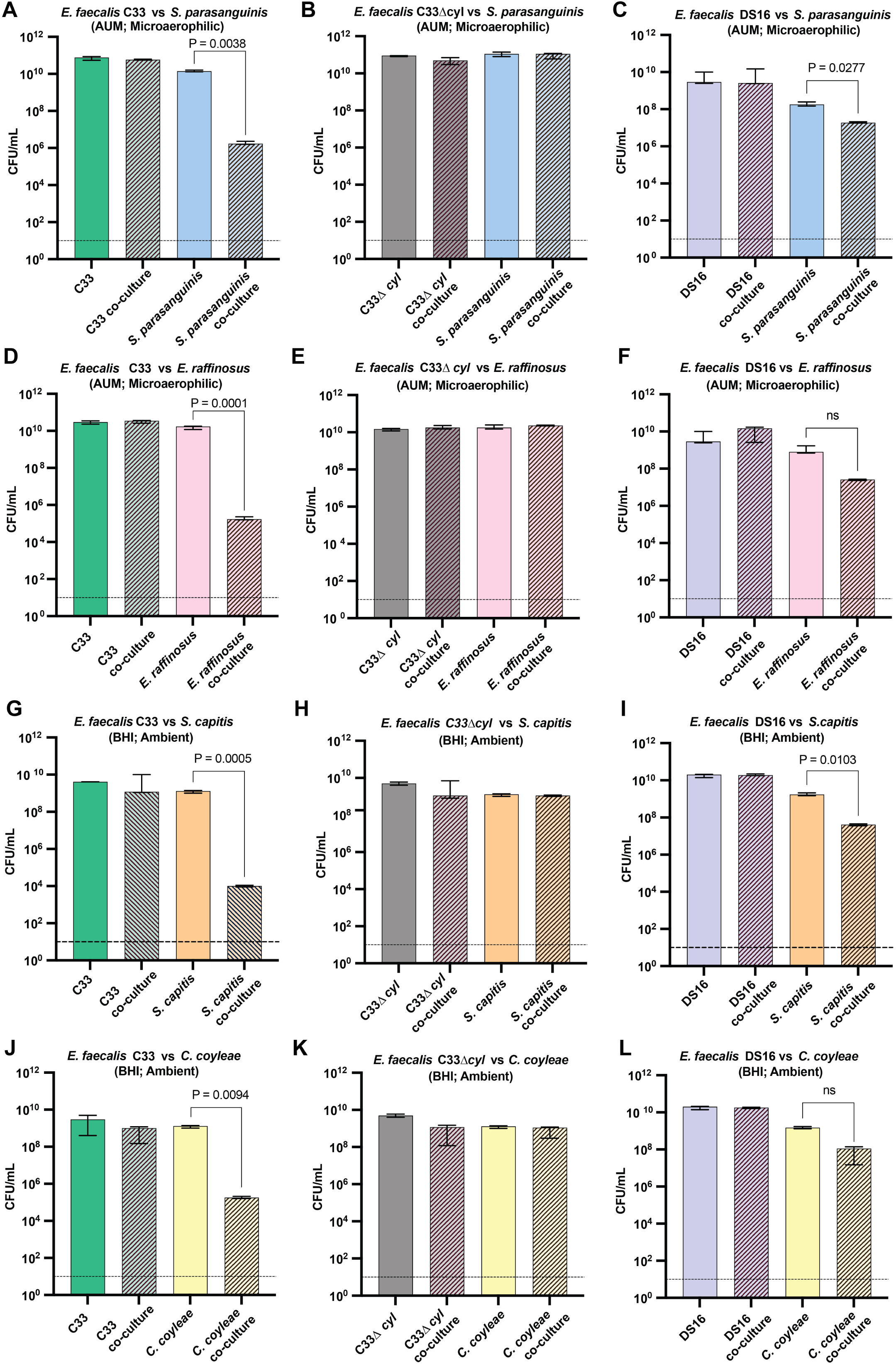
*E. faecalis* C33 cytolysin activity inhibits the urinary microbiota. Quantitative CFU competition assay to assess cytolysin-dependent activity of *E. faecalis* C33 against bacterial species isolated from the same urine sample. Co-cultures of **(A)** *E. faecalis* C33 and *Streptococcus parasanguinis,* **(B)** C33Δ*cyl* and *S. parasanguinis*, **(C)** DS16 **and** *S. parasanguinis* as well as **(D)** *E. faecalis* C33 and *Enterococcus raffinosus*, **(E)** C33Δ*cyl* and *E. raffinosus* and **(F)** DS16 and *E. raffinosus* were grown in AUM under microaerophilic atmosphere. Co-cultures of **(G)** *E. faecalis* C33 and *Staphylococcus capitis*, **(H)** C33Δ*cyl* and *S. capitis,* (**I)** DS16 and *S. capitis,* as well as **(J)** *E. faecalis* C33 and *Corynebacterium coyleae,* **(K)** C33Δ*cyl* and *C. coyleae*, and **(L)** DS16 and *C. coyleae* were grown in BHI under ambient atmosphere. P-values were generated via Welch’s T-test.

Finally, we tested C33 cytolysin activity against commensal species like *Lactobacillus crispatus,* which has been speculated to play a protective role in the urogenital tract, and *Bifidobacterium longum* (43, 46, 47). Using two *L. crispatus* strains previously isolated from the urine of postmenopausal women, we performed bacterial spot inhibition assay in microaerophilic atmosphere on MRS agar to allow robust growth of *L. crispatus* (48). Strong zones of inhibition were observed after spotting overnight cultures of *E. faecalis* C33 onto *L. crispatus* lawns, and this activity was cytolysin dependent (**Table 3**). <u>DS16 also inhibited both *L. crispatus* strains but to a lesser extent than C33. Similar</u> results were observed for *E. faecalis* C33 when spotted on a lawn of *B. longum* urine isolate grown on BHI, with C33 showing moderate inhibition of *B. longum* that was also dependent on cytolysin activity (**Table 3**) (43). Collectively, these results suggest that *E. faecalis* C33 cytolysin activity may provide a competitive advantage against the urinary microbiota, thus possibly supporting niche colonization and persistence in the urinary environment.

**Table 3.** Qualitative spot inhibition assay against *L. crispatus* and *B. longum*.

| Spot | Lawn | Inhibition observed |
| --- | --- | --- |
| *E. faecalis* C33 | *L. crispatus* Lc1226 | +++ |
| *E. faecalis* C33Δcyl | *L. crispatus* Lc1226 | - |
| *E. faecalis* C33ΔcylΔbacL1 | *L. crispatus* Lc1226 | - |
| *E. faecalis* DS16 | *L. crispatus* Lc1226 | + |
| *E. faecalis* C33 | *L. crispatus* Lc1700 | +++ |
| *E. faecalis* C33Δcyl | *L. crispatus* Lc1700 | - |
| *E. faecalis* C33ΔcylΔbacL1 | *L. crispatus* Lc1700 | - |
| *E. faecalis* DS16 | *L. crispatus* Lc1700 | + |
| *E. faecalis* C33 | *B. longum* 1649 | ++ |
| *E. faecalis* C33Δ*cyl* | *B. longum* 1649 | - |
| *E. faecalis* C33Δ*cyl*Δ*bacL1* | *B. longum* 1649 | - |
| *E. faecalis* DS16 | *B. longum* 1649 | + |

+++ denotes strong zone of inhibition, ++ denotes moderate zone of inhibition, + denotes weak zone of inhibition

## DISCUSSION

*E. faecalis* is a known commensal of the human gut microbiome that can also cause opportunistic infections of the urinary tract, blood stream and heart (1, 2). With the ability to form biofilms on urinary catheters and implantable devices, *E. faecalis* is a leading cause of catheter-associated UTIs (CAUTI) and other types of hospital acquired infections (HAI) (5–7). Treatment of enterococcal infections can be complicated by intrinsic as well as acquired resistance to antimicrobials and antibiotics (4). Apart from resistance to antibiotics, *E. faecalis* also possesses various virulence factors that play a role in host colonization and immune evasion (8, 9). For example, *E. faecalis* secretes a lantibiotic called cytolysin that increases the infection severity (9). Cytolysin is encoded on a pheromone-responsive, self-transmissible pAD1 plasmid, that has been implicated in various infections like human bacteremia, rabbit models of endophthalmitis and endocarditis, and mouse peritonitis (10–14). Cytolysin has been reported to lyse multiple eukaryotic cells like human, horse, rabbit and mouse erythrocytes, neutrophils, and macrophages. However, sheep erythrocytes (SEs) have been reported to be resistant to *E. faecalis* cytolysin-induced lysis (22). The presence of phosphatidylcholine (PC) in the susceptible cell membrane, which is lacking in SEs, has been hypothesized to be necessary for cytolysin activity and PC the binding target for *E. faecalis* cytolysin (22). However, the requirement for membrane PC for cytolysin activity has not been biochemically demonstrated and little was known about what phospholipids enterococcal cytolysins may target on host membranes.

Here, we describe a new, gain-of-function cytolysin activity found within select *E. faecalis* isolates that does not require the presence of PC in target membranes for lysis and can efficiently lyse sheep erythrocytes, bladder epithelial cells, and co-isolated species of the urinary microbiota. Characterization of this activity in *E. faecalis* urinary isolate C33 found that hemolysis of sheep erythrocytes but not rabbit erythrocytes increased as atmospheric oxygen decreased, suggesting the gain-of-function cytolysin activity may be regulated by oxygen tension. Although we found that the C33 pAD1-encoded cytolysin operon was necessary for gain-of-function cytolysin activity, transfer of the C33 pAD1 plasmid to an OG1RF recipient strain revealed that the operon and indeed, the entire C33 pAD1 and pAD2 plasmids, were insufficient for SE lysis. These data suggest a chromosomally encoded factor synergizes with the cytolysin operon-encoded lantibiotic to confer the observed gain-of-function activity. Future studies are needed to identify and characterize this chromosomally encoded synergistic factor.

We also demonstrate that PC is indeed required for canonical cytolysin activity by *E. faecalis* blood isolate DS16, which encodes a cytolysin operon identical to the C33 operon also on a pAD1 plasmid. A previous study by Miyazaki *et al*. found that addition of PC or sphingomyelin to horse blood agar plates inhibited *E. faecalis* cytolysin activity against horse erythrocytes (22). These results provided evidence to support the idea that PC or sphingomyelin may be potential binding receptors for the cytolysin lantibiotic but did not conclusively demonstrate that PC was necessary for cytolysin activity. To confirm if canonical DS15 cytolysin requires PC and test if *E. faecalis* C33 gain-of-function is PC-independent, we generated liposomes with varying phospholipid compositions and performed liposome leakage assay. Our results confirm that *E. faecalis* DS16 requires PC for liposome lysis while C33 does not. Interestingly, we found that PE was necessary for lysis by C33 cytolysin but was not required by DS16. This mode of action is similar to another lantibiotic cinnamycin that is produced by *Streptomyces spp*., which also targets PE (49, 50). However, no other annotated lantibiotic biosynthetic gene clusters besides the pAD1-encoded cytolysin operon were found in the *E. faecalis* C33 genome. PE has also been found to enhance the attachment of pathogens like *Helicobacter pylori*, *Haemophilus influenzae*, and both enteropathogenic and enterohemorrhagic *E. coli* to host cells (51). Future studies will explore whether PE mechanistically contributes to *E. faecalis* C33 gain-of-function cytolysin activity.

Because PE is a major phospholipid in the mammalian plasma membrane and cytolysin activity against other eukaryotic cells like retinal cells, leukocytes and intestinal epithelial cells has been reported (21, 52), we performed LDH cytotoxicity assay to test if *E. faecalis* C33 would be more toxic to human 5637 bladder epithelial cells than DS16 (50). Indeed, *E. faecalis* C33 was significantly more toxic (∼80% cytotoxicity) to 5637 bladder epithelial cells than DS16 and \the cytotoxicity was cytolysin-dependent. These data suggest that the *E. faecalis* C33 gain-of-function cytolysin activity may enhance its virulence in the urinary mucosal environment by facilitating the lysis of host epithelial cells. Although 5637 bladder epithelial cells express some markers of differentiated superficial epithelial cells, they are bladder carcinoma cells and not fully representative of the differentiated bladder urothelium. Future studies should employ human bladder organoid or microtissue systems to directly assess the role of this gain-of-function cytolysin activity on *E. faecalis* urovirulence in a fully differentiated uroepithelial environment (53).

*E. faecalis* cytolysin is a eukaryotic toxin as well as a prokaryotic bacteriocin (38, 39). Since *E. faecalis* is an important member of the urinary microbiome and is often part of polymicrobial communities, we asked if C33 cytolysin activity would impact *E. faecalis* community members. Using bacterial strains isolated from the same urine sample as C33 *E. faecalis,* and therefore part of the same microbial community, we found that C33 gain of function cytolysin activity improved *E. faecalis* competition against *Streptococcus parasanguinis*, *Enterococcus raffinosus*, *Staphylococcus capitis* and *Corynebacterium coyleae* as compared to the canonical DS16 cytolysin activity. We also found that *E. faecalis* C33 cytolysin activity inhibited *L. crispatus,* one of the key commensal species in the female urogenital microbiome. Thus, this gain of function cytolysin activity may give urinary *E. faecalis* a competitive advantage over the resident healthy urinary microbiota and within complex polymicrobial communities to establish itself and colonize the urogenital niche.

## MATERIALS AND METHODS

### Bacterial strains and culture conditions

Bacterial strains and plasmids used in the study are listed in **Table 1**. The *E. faecalis* C33 strain was isolated from the urine of a postmenopausal woman with a history of recurrent UTI (43). All *E. faecalis* strains were routinely cultured on Brain-Heart Infusion (BHI) agar, blood agar with 5% sheep blood in ambient and 5% CO_2_ atmospheres, or BD BBL CDC anaerobe 5% sheep blood agar in microaerophilic and anaerobic atmospheres. Liquid cultures were prepared either in BHI broth at 37°C in static conditions, unless otherwise noted. For liposome leakage assays and LDH cytotoxicity assays, *E. faecalis* was grown overnight in RPMI-1640 at 37°C non-shaking, normalized to OD_600_ = 0.1 in fresh media. Antibiotics were used in the following concentrations: Chloramphenicol (Cam), 15 µg/mL; Erythromycin (Erm), 10 µg/mL; Spectinomycin (Spec), 100 µg/mL; Tetracycline (Tet), 10 µg/mL; Rifampicin (Rif), 15 µg/mL.

### Routine molecular biology procedures

Polymerase chain reaction (PCR) was conducted using DreamTaq Master Mix (ThermoFisher) or with Phusion polymerase (New England Biolabs) for fragments ≥ 6 kb in size. Primer sequences for PCR and qRT-PCR are listed in **Table S3**. PCR product purification and plasmid isolation were performed using the EZ-10 Spin Column PCR Products Purification Kit (Bio Basic) and EZ-10 Spin Column Plasmid DNA Miniprep Kit (Bio Basic) as per the manufacturer protocol. Genomic DNA extractions were performed using DNeasy blood and tissue kit (Qiagen). Briefly, 8 mL of bacterial overnight culture was pelleted and incubated lysozyme as per manufacturer instructions coupled with mutanolysin and RNase A treatment. DNA quality was analyzed by agarose gel electrophoresis and quantified by NanoDrop.

### Preparation of competent cells

Chemically competent *Escherichia coli* DH5α cells were used for plasmid propagation. To make DH5α competent cells, overnight cultures of DH5α were sub-cultured to OD_600_ = 0.3 in 2XYT medium at 37°C, and then pelleted at 5000 rpm for 10 min, resuspended in cold competent cell buffer (60 mM CaCl2, 15% glycerol, and 10 mM PIPES at pH 7.0) and incubated on ice for 30 min. Cells were then pelleted and resuspended in cold competent cell buffer and stored at −80°C. For electrocompetent *E. faecalis* cells, overnight cultures were sub-cultured in fresh BHI to OD_600_ = 0.4-0.6, pelleted at 6,000 rpm for 10 min at 4°C then washed with 10% glycerol. The pellet was then resuspended in 500 µL lysozyme solution (10 mM Tris pH 8.0, 20% sucrose, 10 mM EDTA, 50 mM NaCl, 30 µg/ml lysozyme), incubated at 37°C for 20 min and pelleted at 13,000 rpm for 1 min. The pellet was washed four times with 500 µL of electroporation buffer (0.5 M sucrose, 10% glycerol, pH 7.0), aliquoted and stored at −80°C.

### Generation of *E. faecalis* C33 gene deletions and complementations

A markerless gene exchange technique was employed to create deletion and complementation mutants in *E. faecalis*. A derivative of plasmid pLT06 (29), designated as pCOP88 (54), was used for generating mutants. This plasmid contained chloramphenicol (*cat*) selection, p-Cl-phenylalanine counterselection (*pheS*) as well as temperature-sensitive origin (*repA*). To generate an *E. faecalis* C33 cytolysin operon deletion mutant (C33Δ*cyl*), pCOP88 gene sequences *cat*, *pheS* and *oriT* (encoding *repA*) along with 1.0 kb homologous sequences upstream and downstream of the cytolysin operon in *E. faecalis* C33 were amplified by PCR. The resulting fragments were purified, and Gibson assembled into a knock-out plasmid pKO1. The assembled Gibson reaction was transformed into chemically competent *E. coli* DH5α for propagation and plasmid isolation. 1 µg of purified pKO1 plasmid was transformed into 30 µL electrocompetent *E. faecalis* C33 cells by electroporation in a 0.2cm gap width cuvette at 2.5kV using a micropulse electroporator (Biorad). The cells were recovered at 30°C by adding 0.5M sucrose-BHI for 3 hours, followed by plating on BHI + Cam15 plates and incubated for 48 hours. Resulting colonies were streaked on BHI-Cam plates and incubated at the non-permissive temperature of 42°C for 48 hours. Well-isolated colonies were inoculated in 1 mL of BHI broth, incubated at 37°C for 5 hours, and plated on MM9YEG plates supplemented with p-Cl-phenylalanine at 37°C for 24 hours. Bacterial colonies were screened for *cyl* operon deletion by PCR and tested for loss of hemolytic activity. A similar strategy was employed to complement *E. faecalis* C33Δ*cyl* with the cytolysin operon. Briefly, knock-in plasmid pKI, containing the *cat, pheS, oriT*, 1 kb homology regions, and the cytolysin operon sequences was assembled into a plasmid by Gibson. The purified pKI plasmid was transformed into *E. faecalis* C33Δ*cyl* and subjected to the same selection procedure as above. Colonies from MM9YEG plates were screened for presence of the *cyl* operon by PCR and gain of hemolytic function. Additionally, the Bac41 bacteriocin component, *bacL1*, was deleted from both *E. faecalis C33* and C33Δ*cyl* using the same strategy to generate single and double *bacL1* knock-outs.

### Quantitative hemolysis assays

*E. faecalis* strains were grown overnight in RPMI-1640 at 37°C under static conditions. Then, the bacteria were normalized to OD_600_ = 0.05 in RPMI-1640 containing an equal volume (1:1) of either defibrinated sheep or rabbit blood. The cultures were incubated in ambient, 5% CO_2_, microaerophilic and anaerobic atmospheres, at 37°C under static conditions for 24 hours and 48 hours. To quantify the hemolytic activity, cultures were centrifuged for 10 min at 150xg to pellet the erythrocytes, followed by pelleting the bacterial cells for 10 min at 8000xg. Hemoglobin release was measured by determining the absorbance at 540 nm of the supernatant. Percent hemolysis was calculated as follows: ((OD_test_-OD_negative control_)/(OD_positive control_-OD_negative control_))*100. 1% Triton X-100 and RPMI-1640 were used as positive control (100% hemolysis) and negative control respectively. This experiment was performed in biological and technical triplicates.

### Plasmid assembly and alignments

The genomes of urinary *E. faecalis strains* were previously reported in Sharon et al 2023 (26) and were generated via de novo hybrid genome assembly using paired-end Illumina short reads and Oxford Nanopore long reads as previously described (55). *E. faecalis* DS16 (GCA_000395285.1), OG1RF (GCA_000172575.2) and MMH594 (GCA_000394795.1) genomes were obtained from the NCBI database. The closed DS16 pAD1 plasmid was generated by reassembly of the Illumina sequencing reads (SRA SRX158205) using SPAdes v3.15.2. Assembled contigs were screened for pAD1-like plasmid sequences using blastn from NCBI BLAST+ v2.12.0+ against pAD1 reference plasmid sequences from *E. faecalis* strains C33 and C130. Candidate plasmid contigs were assessed based on BLAST similarity, contig length, and assembly coverage. The pAD1 contig was circularized in Geneious Prime 2024.0.3 and used for downstream comparison. Comparative plasmid alignment and visualization were conducted using BLAST Ring Image Generator v0.95 with blastn at default thresholds (56). The C33 pAD1 plasmid sequence was used as the reference. Pairwise nucleotide similarity and plasmid coverage were calculated from blastn high-scoring segment pairs by merging overlapping aligned intervals and dividing the nonredundant aligned length by the total plasmid length.

### Cytolysin operon sequence aligment

The cytolysin operon regions from the pAD1 plasmid of *E. faecalis* DS16 and the corresponding cytolysin operon region from *E. faecalis* C33 were extracted in Geneious Prime 2024.0.3 and exported in GenBank format. Alignment of the cytolysin operons was conducted using Easyfig v2.2.5 with blastn at default parameters. Pairwise nucleotide identity and coverage across the operon were confirmed from the blastn alignment output.

### RNA extraction, cDNA synthesis, and qRT-PCR

For RNA extractions, overnight cultures were sub-cultured to an OD_600_ of 0.1 in RPMI-1640 containing equal volume sheep blood, followed by incubation in either anaerobic or microaerophilic atmosphere for 4 hours until mid-log phase (OD_600_=0.4-0.6) under static conditions. Total RNA was extracted from cultures using the Qiagen RNeasy Plus kit following the manufacturer’s instructions. RNA concentration and purity were analyzed by NanoDrop (Thermo). cDNA synthesis was performed on RNA samples with A260/280 ratios between 2.00 and 2.20 using the Qscript cDNA kit (Quantabio). qRT-PCR was performed using PerfeCTa SYBR Green FastMix (Quantabio). *adk* gene was used as the housekeeping gene to normalize the expression of gene of interest. Relative gene expression was calculated by the delta-delta Ct method (2^-ΔΔCt^). This experiment was performed in biological and technical triplicate.

### Conjugation assay

Mating assays with OG1RF were performed with an OG1RF strain encoding *E. faecalis* C33 *cylA* & *cylI* genes on a cCF10 pheromone inducible plasmid. After overnight growth, the cultures were sub-cultured statically in BHI at 37°C until OD_600_ ∼0.5. For OG1RF:pCIE-*cylAI,* 10 ng/ml of cCF10 pheromone was added to the sub-culture. Donor (*E. faecalis* C33) and recipient cultures (*E. faecalis* OG1RF:pCIE-*cylAI*) were mixed in a 1:9 ratio and then pelleted at 6,000xg for 2 minutes. The pellet was reconstituted in 100 µL and then spotted on BHI agar and incubated at 37°C overnight. The cells were then scraped into 1 mL of 1X Phosphate-Buffered Saline (PBS) with 2 mM EDTA, serially diluted, and plated on BHI agar supplemented with Rif15 to select for OG1RF transconjugants. Transconjugants were screened for gain of either pAD1 or pAD2 or both plasmids by PCR targeting pAD1-*cylB*, pAD2-*bacL1* and OG1RF-*rhaB* genes.

### Liposome leakage assay

Phospholipids used in the assay were purchased from Avanti Polar Lipids. Liposomes were prepared using the Avanti Research Mini-Extruder extrusion technique. Briefly, the lipid mix was dried under nitrogen stream, followed by drying under vacuum overnight. Then, lipids were resuspended in RPMI-1640 to set the final liposome concentration at 5 mM. 5(6)-carboxyfluorescein fluorescent dye prepared in ethanol was added at a concentration of 130 µM. Small unilamellar vesicles were generated by three freeze thaw cycles of the lipids in liquid nitrogen, followed by successively extruding 11 times each through polycarbonate membranes with decreasing pore sizes (1 µm, 0.4 µm and 0.2 µm) using a 1 ml air-tight syringe system. Liposomes were then harvested by ultracentrifugation at 180000 x g, for 30 min at 4°C. The liposomal pellets were washed twice in RPMI-1640 and then resuspended in RPMI-1640 to a final concentration of 5 mM. Liposome size distribution and polydispersity was determined by UV-visible dynamic light scattering (633 nm laser wavelength, scattering angle of 175°, medium refractive index of 1.33 and material refractive index of 1.51). Liposomes were diluted 1:5 in RPMI-1640 and incubated with *E. faecalis* for 4 hours. Liposomes treated with 1% Triton X-100 for 30 min served as positive/high controls for liposome lysis and untreated liposomes were negative/low controls. After incubation, samples were pelleted by ultracentrifugation at 180000 x g, 30 min at 4°C. The supernatant was transferred to a 96-well black, clear-bottom microplate with and fluorescence measured at 484/530 nm (excitation/emission). Percentage lysis was calculated as follows: ((Fluorescence of *E. faecalis* treated liposome well – fluorescence of low control well)/ (Fluorescence of high control– fluorescence of low control wells))*100. All experiments were performed in biological triplicates with three technical replicates for each.

### LDH release assay

LDH Cytotoxicity Detection Kit (Takara) was used to measure the LDH release. Urinary bladder epithelial 5637 cells (ATCC no. HTB-9) were cultured in RPMI-1640 (Sigma) supplemented with 10% Fetal Bovine Serum (FBS) (ThermoFisher), 1% penicillin-streptomycin (Gibco), and 1% glutamax (Gibco) at 37°C and in 5% CO_2_ atmosphere overnight. Cells were seeded into 24 well plates and after reaching 80-90% confluency, cells were washed in colorless RPMI-1640 without antibiotics, then infected with *E. faecalis* strains at a multiplicity of infection (MOI) of 10 and incubated at 37°C with 5% CO_2_ for 4 hours and 16 hours. 5637 cells treated with 1% Triton X-100 for 10 minutes served as positive/high controls for maximal LDH release while untreated cells served as low controls. Following incubation, 200 µL of supernatant was transferred from each well to 96-well plate and centrifuged at 1000xg for 5 min to pellet the cell debris. 100 µL of cell free supernatant was transferred to a new 96-well plate. To measure LDH release, 100 µL of the reaction mixture (diaphorase/NAD+ catalyst and INT/Na-lactate dye solution) was added to the wells and absorbance was measured at 490 nm using a microplate reader (BioTek Synergy H1). Cytotoxicity was expressed as a percentage of maximal LDH release using the following formula: % Lysis = ((OD value of *E. faecalis* treated cells – OD of low control)/ (OD of high control – OD of low control))*100. The experiment was performed in biological triplicates with three technical replicates for each.

### Spot inhibition assay

*B. longum* was cultured in BHI under 5% CO_2_ and *L. crispatus* were grown in MRS in microaerophilic atmosphere (48). Overnight cultures of *E. faecalis* grown in BHI or De Man, Rogosa and Sharpe (MRS) were normalized to OD600 = 0.05 in fresh medium. After normalization, the culture was swabbed onto BHI agar for testing *Bifidobacterium longum* or MRS agar for testing *Lactobacillus crispatus* to form a lawn. 10 µL of the tester strain (*B. longum* or *L. crispatus*) was spotted onto the lawn and the spots were allowed to dry. The plates were incubated in 5% CO_2_ or microaerophilic atmosphere overnight. Plates were imaged the following day for zone of inhibition.

### Quantitative competition assay

For *S. parasanguinis* and *E. raffinosus* assays, bacteria were grown in AUM^Y^ overnight, followed by normalization to OD600 = 0.05 and were either incubated individually or co-cultured with *E. faecalis* strains in AUM^Y^ at microaerophilic atmosphere overnight in static conditions. Following overnight incubation, the cultures were serially diluted in sterile 1X PBS and 10 µl of the serial dilution was spotted onto BHI agar supplemented with antibiotics for selection. Tet10 or Erm10 was used for selecting urinary *E. faecalis* strains, DS16 and MMH594. Rif15 was used for OG1RF. Additionally, if no differential intrinsic antibiotic resistance was found for selection, the strains were transformed with pKH12 (Cam15) or pAT28 (Spec100) plasmids to confer antibiotic resistance, if required. A similar protocol was followed for *S. capitis* and *C. coyleae* except they were cultivated aerobically in BHI instead as AUM did not support growth. Following overnight incubation of the *S. capitis*-*E. faecalis* C33 and *C. coyleae*-*E. faecalis* C33 co-cultures, they were serially diluted, plated onto chromogenic agar (ChromAgar Orientation, BD) to enable differential identification, and incubated overnight at 37°C in an ambient atmosphere. CFU/mL for each strain was calculated using the following formula: (Number of colonies) / (dilution factor * volume of culture spotted on the plate). All the experiments were performed in biological triplicate and technical triplicate.

## ACKNOWLEDGEMENTS

This work was funded by the Welch Foundation, award numbers AT-2030-20200401 to N.J.D. and AT-2073-20240404 to G.M., by the National Institutes of Health, award number R01DK131267 to N.J.D. and R35GM128704 to G.M., and by the Felecia and John Cain Distinguished Chair in Women’s Health, held by P.E.Z.

## AUTHOR CONTRIBUTIONS

Conceptualization: N.V.H. and N.J.D.; Data curation: N.V.H., H.G., I.N.H., I.O.S., N.V.S., U.B.; Formal analysis: N.V.H., H.G., I.N.H., I.O.S., N.V.S.; Investigation: N.V.H.; H.G., I.N.H., I.O.S.; Methodology: N.V.H., H.G., I.O.S., N.V.S., G.M., N.J.D.; Funding acquisition: P.E.Z. and N.J.D.; Project administration: N.J.D. and G.M.; Resources: P.E.Z., G.M., N.J.D.; Supervision: G.M. and N.J.D.; Validation: N.V.H. and N.J.D.; Visualization: N.V.H., I.O.S, N.J.D.; Writing original draft: N.V.H., I.O.S and N.J.D.

