## Supplemental material for "Gain of Function Cytolysin Variant Expressed by Clinically Isolated Urinary *Enterococcus faecalis*"

Hulyalkar et al.

Tables S1-S3

Figures S1-S6

Table S1. Additional liposome compositions used for liposome leakage assay

| **Phospholipids** | **Rabbit Erythrocyte (RE)-like (%)** | **RE-like w/o PS (%)** | **RE-like w/o PI (%)** | **RE-like w/o**  **SM (%)** |
| --- | --- | --- | --- | --- |
| Phosphatidylethanolamine (PE) | 32 | 35 | 32.5 | 37 |
| Phosphatidylserine (PS) | 12 | 0 | 12.5 | 17 |
| Phosphatidylinositol (PI) | 2 | 5 | 0 | 7 |
| Phosphatidylcholine (PC) | 34 | 37 | 34.5 | 39 |
| Sphingomyelin (SM) | 20 | 23 | 20.5 | 0 |

Table S2. Artificial Urine Medium supplemented with 0.3% Yeast extract (AUMY) for growth of *E. faecalis.*

| **Component** | **Quantity** | **Unit (per liter)** |
| --- | --- | --- |
| Peptone L37 | 1 | g |
| Yeast Extract | 0.005 | g |
| Sodium Bicarbonate | 2.1 | g |
| Sodium Chloride | 5.2 | g |
| Disodium Sulfate Decahydrate | 3.2 | g |
| Potassium Dihydrogen Phosphate | 0.95 | g |
| Dipotassium Hydrogen Phosphate | 1.2 | g |
| Ammonium Chloride | 1.3 | g |
| L-Lactic Acid | 0.1 | g |
| Urea | 10 | g |
| Citric Acid | 0.4 | g |
| Creatinine | 0.8 | g |
| Iron (II) Sulfate Heptahydrate | 0.0012 | g |
| Uric Acid | 0.07 | g |
| Calcium Chloride | 0.37 | g |
| Magnesium Sulfate | 0.49 | g |
| Yeast Extract^**^ (For AUM^Y^) | 3 | g |

TABLE S3. Polymerase Chain Reaction (PCR) & Quantitative real-time PCR (qPCR) primer sequences used in this study.

| **PCR primers** | | |
| --- | --- | --- |
| **Primer name** | **Description** | **Primer Sequence (5’-3’)** |
| C33_Arm1_F | Knock-out of cytolysin (*cyl*) operon in C33 | ACCTCAAATGGTTCGCTGGGTTTA  TCGACCCTAATTTTTCTAGCTACAGTATTTTTG |
| C33_Arm1_R | Knock-out of *cyl* operon in C33 | GTCTAGCTGGTTTCTTATCAAA GATTTTTAAATTTCACCACAGG |
| C33_Arm2_F | Knock-out of *cyl* operon in C33 | AAATTTAAAAATCTTTGATAA  GAAACCAGCTAGACTG |
| C33_Arm2_R | Knock-out of *cyl* operon in C33 | CTAAAACGTCCTATTCAAAGAAACGCT GCACTATGTTTAAGTATTAACAATCTTC |
| *ori*_F | Ori for knock-out/in plasmid | TGCAGCGTTTCTTTGAATAG |
| *ori*_R | Ori for knock-out/in plasmid | GCTTTGCAAAGTCTGAAAAC |
| *pheS-cat*_F | Pos/neg selection for knock  out/in plasmid | TGCCACCTTCGTTTTCAGACTTTGCAA AGCCAAGTTAAGGGATGCAGTTTAAA  AATG |
| *pheS-cat*_R | Pos/neg selection for knock  out/in plasmid | GGTCGATAAACCCAGCGA |
| C33_Screen_F | Verify *cyl* operon deletion | CAGTTATGATTAAAGTATAAAACAAC |
| C33_Screen_R | Verify *cyl* operon deletion | CGTATGCCCAACTTGAATG |
| C33_*cyl_*operon_F | Verify *cyl* operon deletion | TTATTCAGGTTGCCATTGAAAAATATC |
| C33_*cyl_*operon_R | Verify *cyl* operon deletion | TCAATTTTGAACCCATTGTTTTTCTG |
| C33_*cylB*_F | Verify *cyl* operon  deletion/complementation | ATGAAAAGATTGAAGTACGTTGC |
| C33_*cylB*_R | Verify *cyl* operon  deletion/complementation | TCATTTTGTACCTCCAAATTGTTG |
| C33_*cylM*_F | Verify *cyl* operon  deletion/complementation | GGAAGATAATTTAATAAATGTTTTAAGC |
| C33_*cylM*_R | Verify *cyl* operon deletion/complementation | CTTTAGAGGTTCATTTTTTTTAAAAG |
| C33_*cylA*_F | Verify *cyl* operon deletion/complementation | ATGAAAAAAAGAGGATTGAC |
| C33_*cylA*_R | Verify *cyl* operon deletion/complementation | CTATTTTATTCTCACCTCTTTG |

| **Primer name** | **Description** | **Primer Sequence (5’-3’)** |
| --- | --- | --- |
| Arm1_F | Knock-in *cyl* operon in C33Δ*cyl* | ACCTCAAATGGTTCGCTGGGTTTA TCGACCGAGCCTTCGATTGCCGTC |
| Arm1_*cyl_*R | Knock-in *cyl* operon in C33Δ*cyl* | ATGGCAACCTGAATAAATAGG  AGCGGTCCTTATTTTTTATTTCTGG |
| Arm1_*cyl*_F | Knock-in *cyl* operon in C33Δ*cyl* | GAAATAAAAAATAAGGACCGCTCCTA  TTATTCAGGTTGCCAT TGAAAAATATC |
| *cyl*_Arm2_R | Knock-in *cyl* operon in C33Δ*cyl* | GTTCTTGATTTAAAATCGATAATGAAA TACACGTCATTCAATTTTGAACCCATT  GTTTTTCTG |
| *cyl*_Arm2_F | Knock-in *cyl* operon in C33Δ*cyl* | CAGAAAAACAATGGGTTCAAAATTGA ATGACGTGTATTTCATTATCGATTTT  AAATC |
| Arm2_R | Knock-in *cyl* operon in C33Δ*cyl* | CTAAAACGTCCTATTCAAAGAAACGC TGCAAAAGGAACCGTCCTTATTTTTT  ATTTG |
| BacL1_Arm1_F | Knock-out of *bacL1* gene in C33 | ACCTCAAATGGTTCGCTGGGTTTATCG  ACCTCGTTTTCTTTTTAACTGTTCGAAA G |
| BacL1_Arm1_R | Knock-out of *bacL1* gene in C33 | CTAATAACTAGTGTTTAAAGCA  ATATAATTTGTAAAACTTCACC TCATATTTTTTGTCG |
| BacL1_Arm2_F | Knock-out of *bacL1* gene in C33 | GTCGCGACAAAAAATATGAGGTGAAGTTTTACAAATTATATTGCTTTAAAC  ACTAG |
| BacL1_Arm2_R | Knock-out of *bacL1* gene in C33 | CTAAAACGTCCTATTCAAAGAAACGC  TGCATTCCTATTCTTGAT TGAAAGTC |
| BacL1_F | Verify *bacL1* deletion in C33 | ATGAATTACAGTCAAAAAGCAATCGATC |
| BacL1_R | Verify *bacL1* deletion in C33 | CTAATTAAAGAATCCTTTGCCCCATAC |
| *cylAI*_F | Expression of C33 *cylA* & *cylI*  genes in pCIE vector | CTAGGGATCCATGAAAAAAAGAGGATTGACTTATATTTTAATC |
| *cylAI*_R | Expression of C33 *cylA* & *cylI*  genes in pCIE vector | CTAGAAGCTTTCAATTTTGAACCCATTGTTTTTCTGC |
| OG1RF_*rhaB*_F | OG1RF-specific rhaB primer | GTGAAAAAAGAACAAGTTGAGAAAC |
| OG1RF_*rhaB*_R | OG1RF-specific rhaB primer | TCATAATGAGCTCCTTTTCATAAATG |

| **qPCR Primers** | | |
| --- | --- | --- |
| **Primer name** | **Description** | **Primer Sequence (5’-3’)** |
| *adk*_F | Housekeeping gene | ATGTGGCGCAACGTATCA |
| *adk*_R | Housekeeping gene | CCGTTTCGGGCTTATCATCT |
| *clpX_*F | Housekeeping gene | CTCCACCAACGATGAAGAGAAC |
| *clpX_*R | Housekeeping gene | TGCAAGTGTACCACCACAAG |
| *cylLL_*F | Expression of cylL_L_ gene | CTGTTGCGGCGACAGCT |
| *cylLL_*R | Expression of cylL_L_ gene | CCACCAACCCAGCCACAA |
| *cylLS_*F | Expression of cylL_S_ gene | GCTAAATAAGGAAAATCAAGAAAA  CTATTACTC |
| *cylLS_*R | Expression of cylL_S_ gene | CAAAAGAAGGACCAACAAGTTCTAATT |
| *cylM_*F | Expression of cylM gene | GTTGGTGTAAGGGAACTGTAGG |
| *cylM_*R | Expression of cylM gene | TCCATGGCACAGACAATCTTTA |
| *cylB_*F | Expression of cylB gene | GTGAATGTGCTTTGGCATGTAT |
| *cylB_R* | Expression of cylB gene | CTTGATAGTTAGTCCTCCTTTGGG |
| *cylA_*F | Expression of cylA gene | CCTACATCGCTAGTTTCTCCAC |
| *cylA_R* | Expression of cylA gene | TGCTAAAGCTGCGCTTACT |
| *cylR2_*F | Expression of cylR2 gene | TCTTCTAGTGGAGTATTCAGGTAGT |
| *cylR2_R* | Expression of cylR2 gene | GCCAAAGTGAATTAGCTGCTTTAT |
| *bacL1_*F | Expression of C33 *bacL1* | AAGCTGGTCAGGCCATTATC |
| *bacL1_*F | Expression of C33 *bacL1* | AAGCTGGTCAGGCCATTATC |
| *rhaB_*F | Expression of OG1RF *rhaB* | GCTTCTAGCGGTCGCATAAT |
| *rhaB_*R | Expression of OG1RF *rhaB* | CCAACGCTGATACCCATCTT |

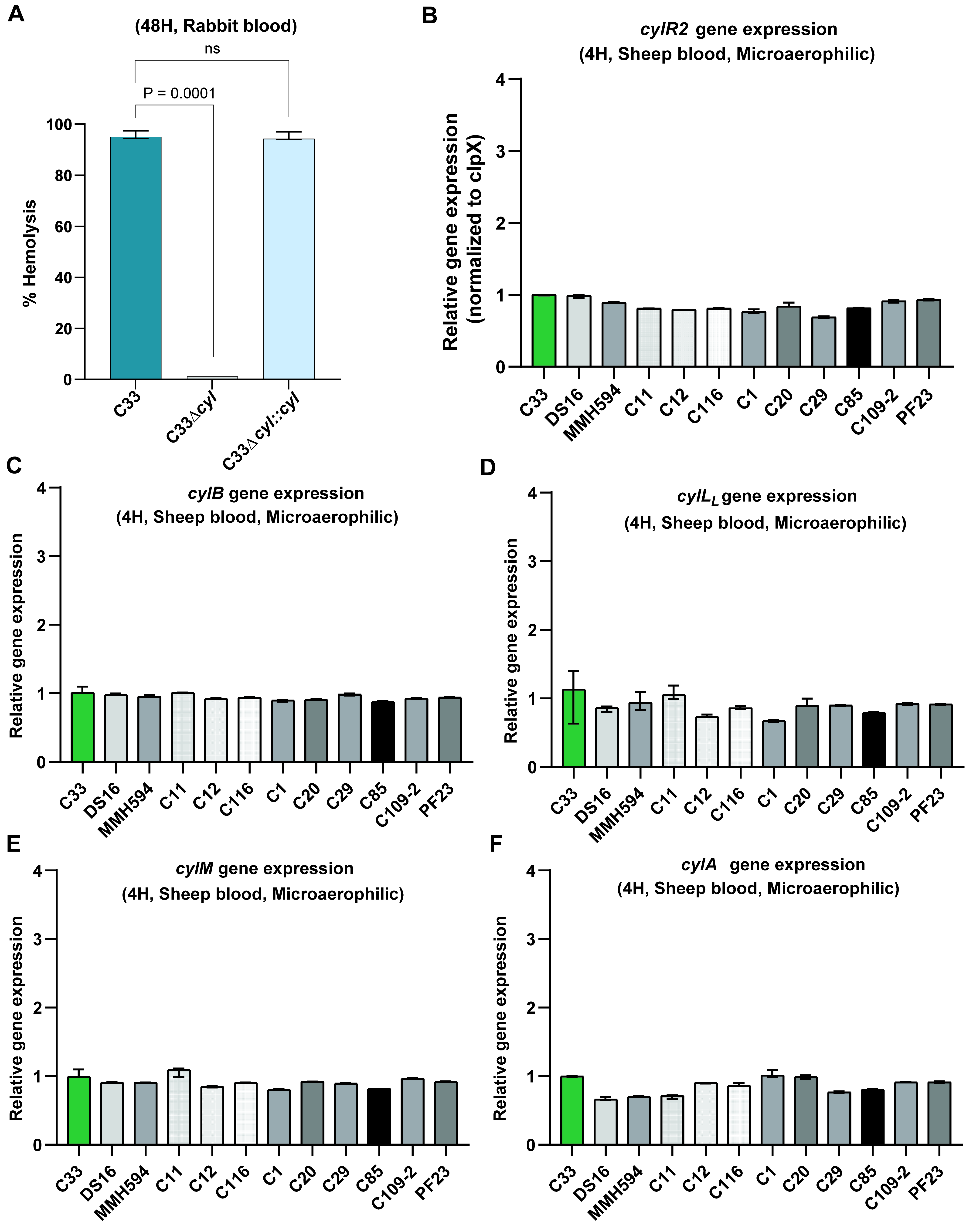

**Figure S1. *E. faecalis* C33 lysis of rabbit erythrocytes is cytolysin-dependent and no difference observed in cytolysin genes expression between *E. faecalis* strains.** Quantitative hemolysis assays to test activity of **(A)** C33, C33Δ*cyl* & C33Δ*cyl*::*cyl* against rabbit erythrocytes. Quantitative Real-time PCR to measure cytolysin genes expression in sheep blood under microaerophilic atmosphere in urinary *E. faecalis* strains **(B)** *cylR2* **(C)** *cylB* **(D)** *cylL_L_* **(E)** *cylM* **(F)** *cylA*. P-values generated via the Brown-Forsythe and Welch ANOVA test.

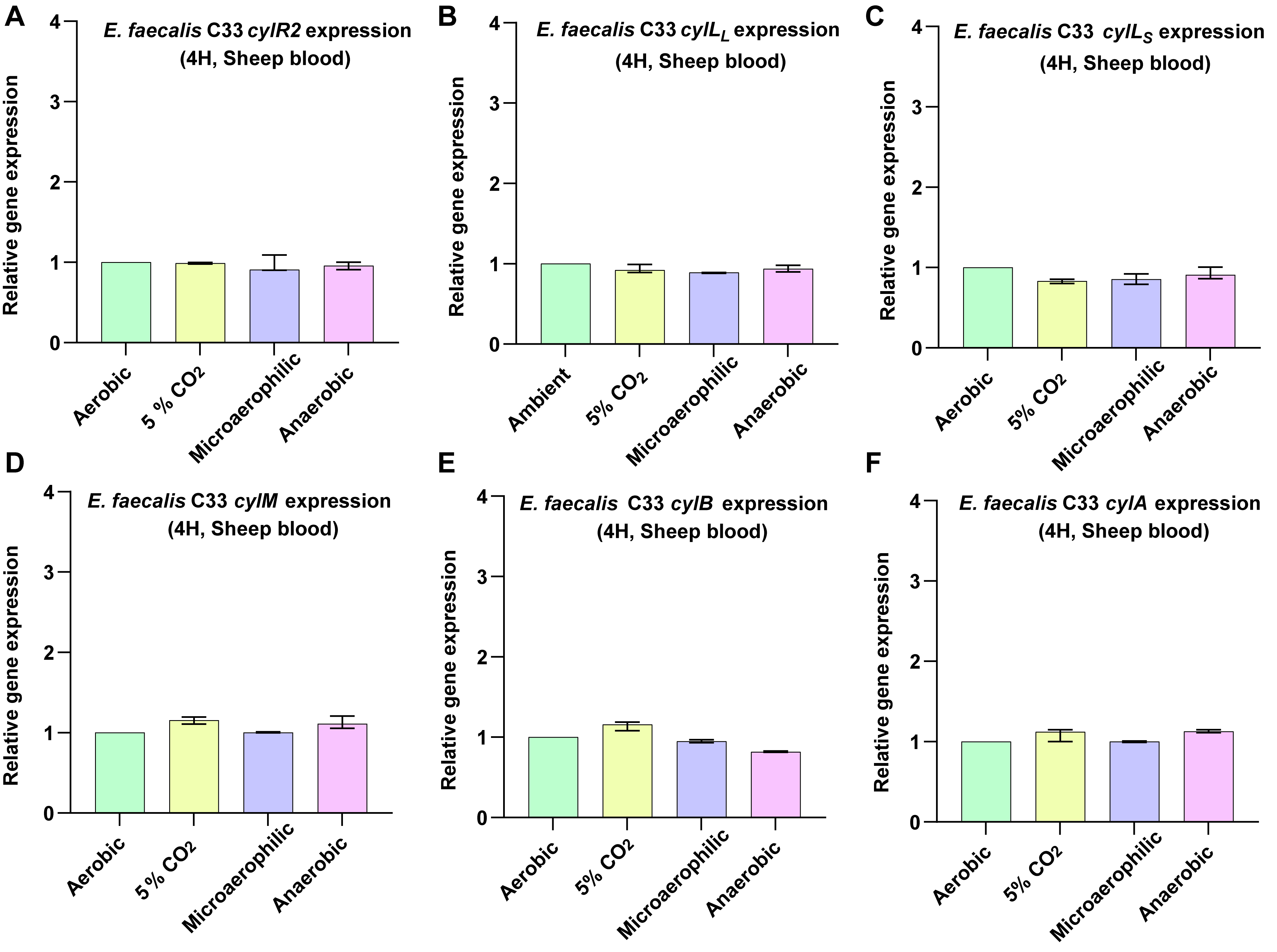

**Figure S2. Cytolysin operon gene expression between different atmospheres in *E. faecalis* C33.** Quantitative Real-time PCR to measure cytolysin genes expression in sheep blood under aerobic, 5% CO2, microaerophilic & anaerobic atmospheres in urinary *E. faecalis* C33 **(A)** *cylR2* **(B)** *cylL_L_* **(C)** *cylL_S_* **(D)** *cylM* **(E)** *cylB* **(F)** *cylA*.

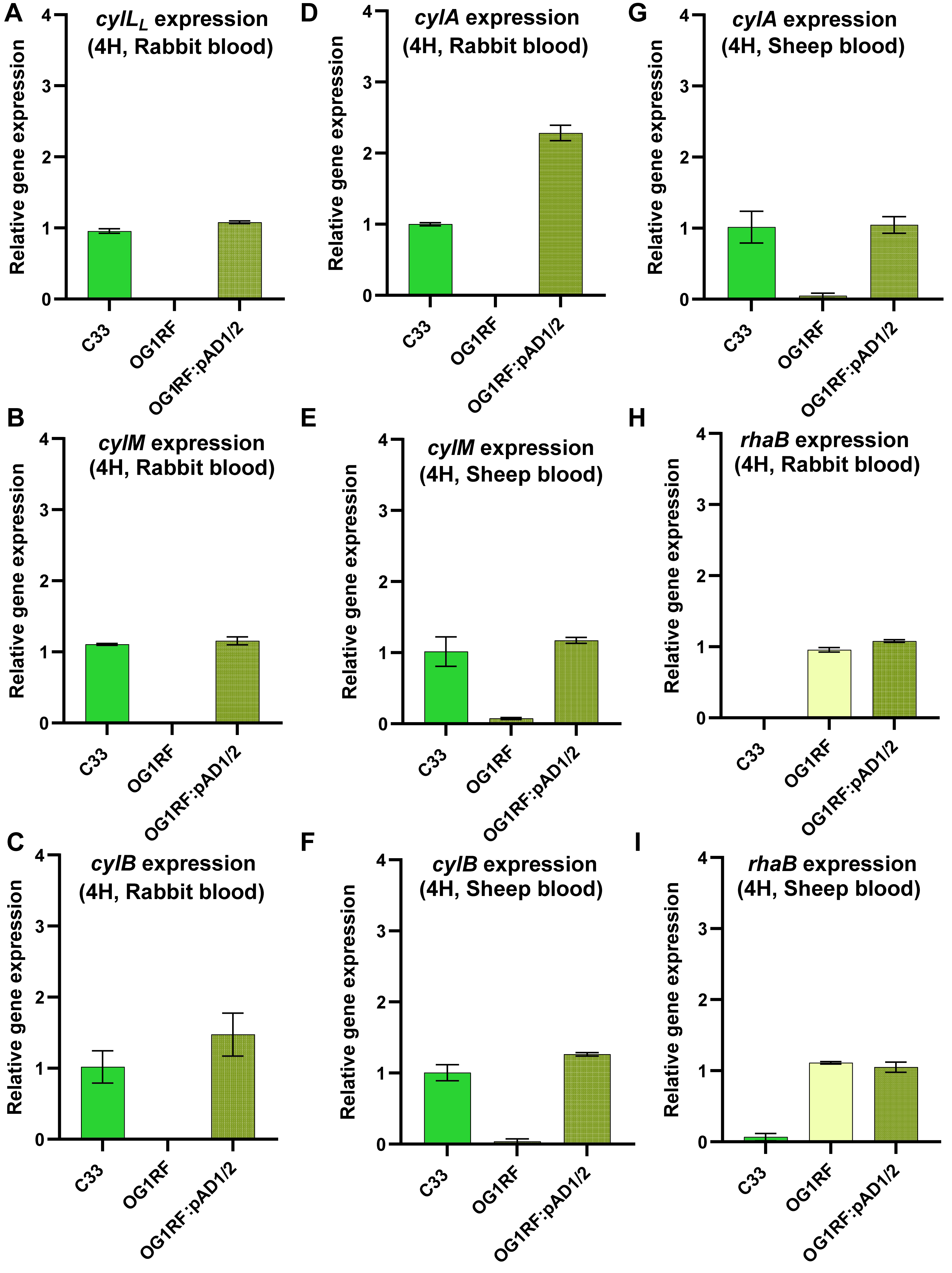

**Figure S3. Transconjugant OG1RF:pAD1/2 expresses cytolysin genes.** Quantitative Real-time PCR to measure cytolysin genes & OG1RF-specific rhaB gene expression under microaerophilic atmosphere in *E. faecalis* C33, OG1RF & OG1RF:pAD1/2 strains; in sheep blood **(A)** *cylL_L_* in rabbit blood **(B)** *cylM* in rabbit blood **(C)** *cylB* in rabbit blood **(D)** *cylA* in rabbit blood **(E)** *cylM* in sheep blood **(F)** *cylB* in sheep blood **(G)** *cylA* in sheep blood **(H)** *rhaB* in rabbit blood **(I)** *rhaB* in sheep blood.

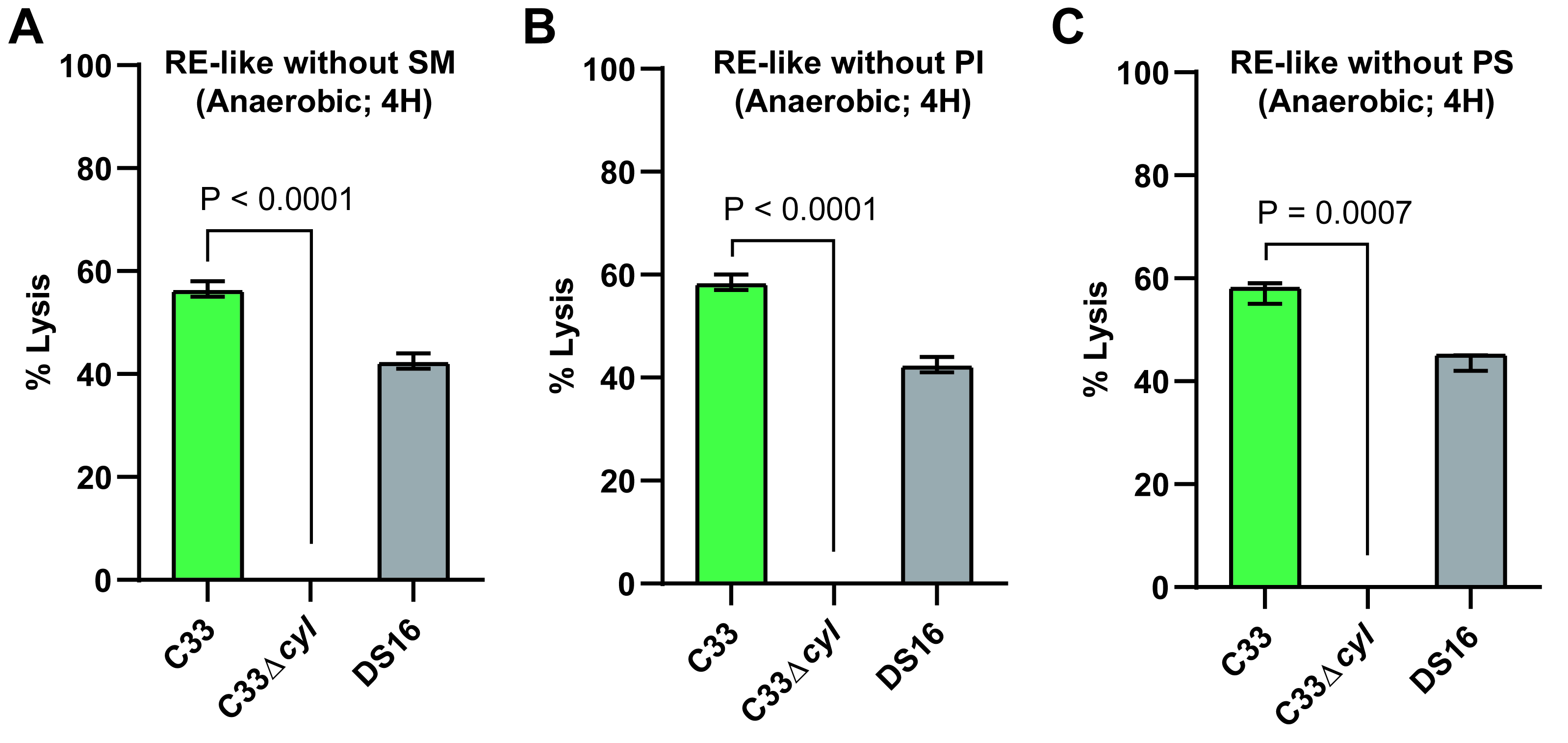

**Figure S4. *E. faecalis* C33 liposome lysis was independent of sphingomyelin phosphatidylinositol & phosphatidylserine lipids.** Liposome leakage assay to measure lysis by C33, C33Δ*cyl* & DS16 in the absence of **(A)** Sphingomyelin **(B)** Phosphatidylinositol **(C)** Phosphatidylserine. P-values generated via the Brown-Forsythe and Welch ANOVA test.

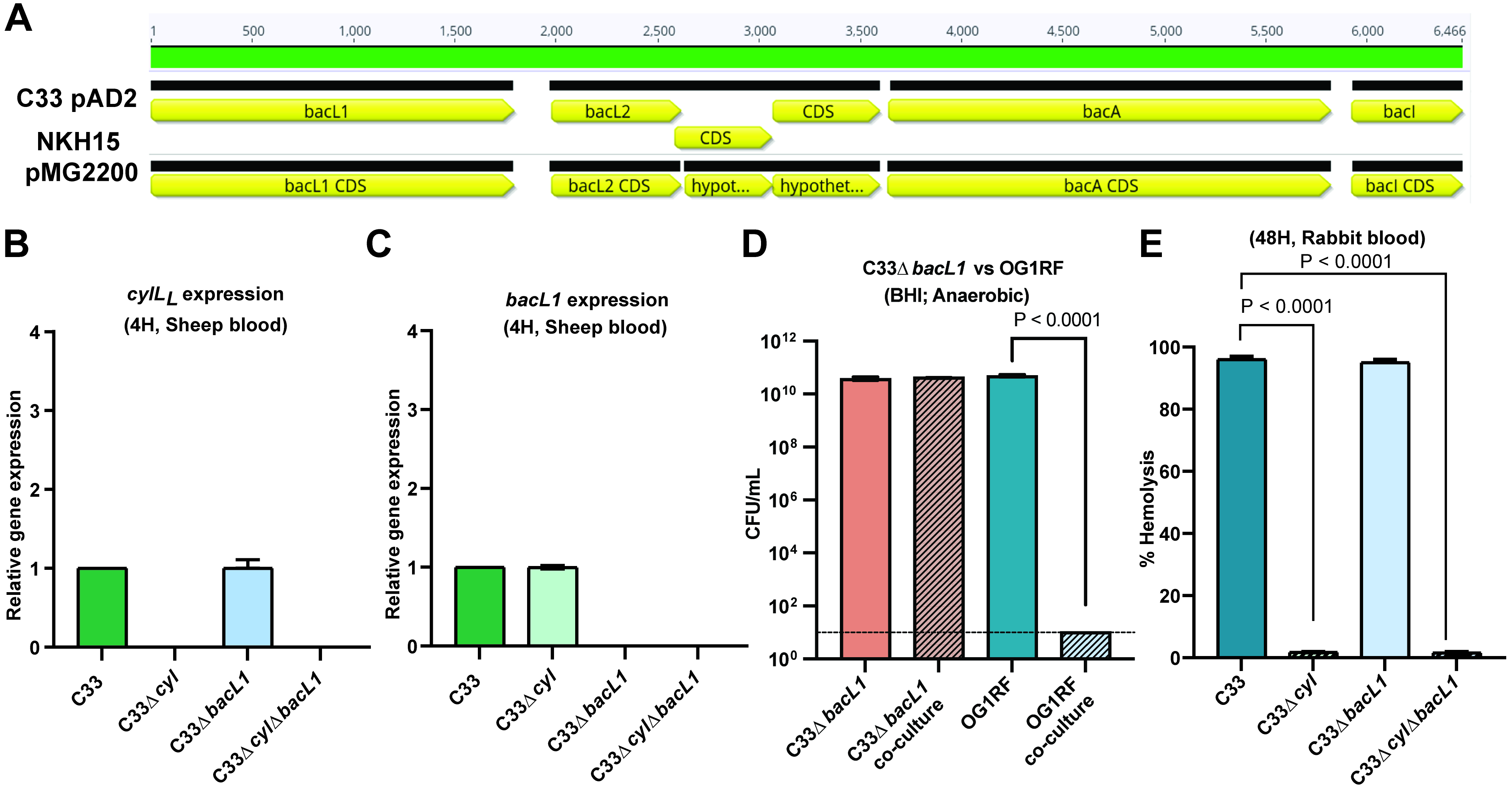

**Figure S5. C33 cytolysin and Bac41 both inhibited *E. faecalis* OG1RF. (A)** Nucleotide sequence alignment of *bacL1* operon genes from C33 and NKH15 strains. Quantitative real-time PCR gene expression of **(B)** *cylL_L_* and **(C)** *bacL1* in C33, C33Δ*cyl*, C33Δ*bacL1* and C33Δ*cyl*Δ*bacL1.* **(D)** CFU competition assay with C33Δ*bacL1* and OG1RF in BHI under anaerobic atmosphere. P-values generated via Welch’s T-test. **(E)** Quantitative hemolysis assay for C33, C33Δ*cyl*, C33Δ*bacL1* and C33Δ*cyl*Δ*bacL1* in rabbit blood. P-values generated via the Brown-Forsythe and Welch ANOVA test.

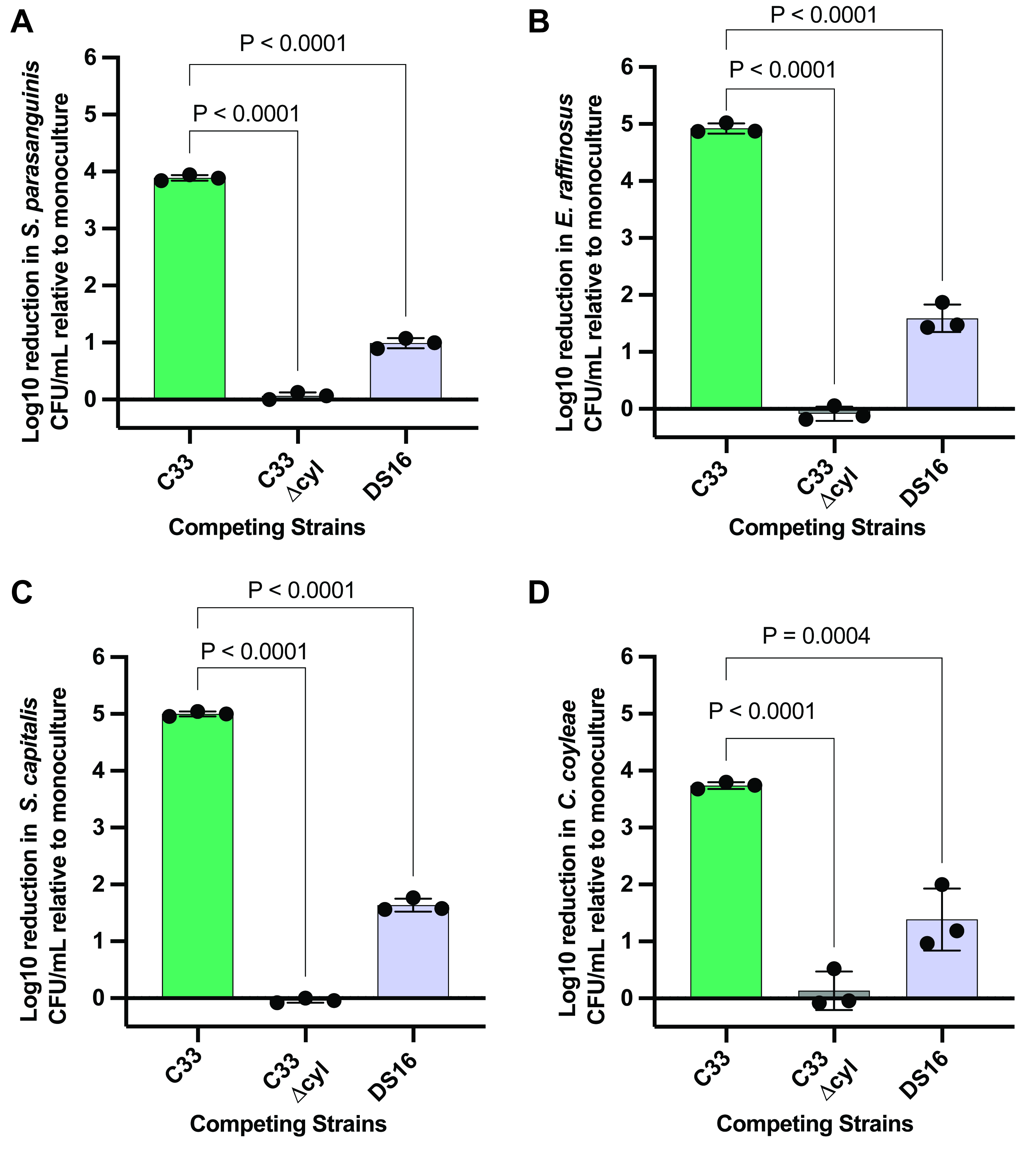

**Figure S6. *E. faecalis* C33 displays greater inhibition of the urinary microbiota during co-culture.** Log₁₀ reduction in CFUs relative to monoculture following co-culture with *E. faecalis* C33, C33Δcyl, or DS16 against **(A)** *S. parasanguinis* **(B)** *E. raffinosus* **(C)** *S. capitis*, and **(D)** *C. coyleae*. Statistical significance was determined by one-way ANOVA with multiple comparisons to the C33 group. Data represent three biological replicates.
